# 1,3-1,6 β-glucans reduce aging hallmarks in multiple organs and rapidly induce mitochondrial biogenesis and autophagy via direct effect on the killifish brain and human neurons

**DOI:** 10.64898/2026.05.15.725450

**Authors:** L. Brogi, B. Fronte, F. Tonelli, M. Marchese, F. Cremisi, A. Cellerino

## Abstract

The short-lived annual fish *Nothobranchius furzeri* (*Nfu*) is a powerful vertebrate model for aging research due to its rapid lifespan and accelerated development of age-associated phenotypes, including gliosis and lipofuscin accumulation. Here, we investigated the effects of dietary 1,3-1,6 β-glucans (BGs), natural polysaccharides derived from *Saccharomyces cerevisiae*, on aging-related processes across multiple tissues, with particular focus on the brain.

Chronic treatment with BG-fortified food reduced several hallmarks of aging in multiple organs. Mechanistically, BG treatment modulated pathways associated with autophagy, lysosomal function, protein oxidation, and inflammation. Both acute and chronic BG administration increased autophagic activity in the aging brain, although lipofuscin accumulation was not affected.

To assess whether BGs act directly on neural tissue, we established an *ex-vivo Nfu* brain culture system that recapitulates the age-dependent decline in autophagy observed in vivo. In this model, acute BG treatment restored impaired autophagy and promoted mitochondrial and lysosomal biogenesis in aged brains. Proteomic analyses revealed increased mitochondrial respiration and modulation of V-ATPase components involved in autophagosome acidification. Depletion of microglia reduced but not eliminated this effect, suggesting direct action of BGs on neurons.

To verify the validity of these findings in humans, we performed BG treatment in human iPSC-derived neurons under conditions of impaired autophagy and found an increase in survival.

Together, these findings identify β-glucans as modulators of autophagy, mitochondrial function, and inflammation, highlighting their potential to promote healthy aging.

## INTRODUCTION

Aging is a universal and multifactorial process that is the principal risk factor for most chronic and degenerative diseases. It is associated with a series of canonical cellular and molecular hallmarks, among which systemic chronic low-grade inflammation (inflammaging) and an autophagy decline that naturally occurs with age (López-Otín et al., 2013; Schmauck-Medina et al., 2022; Aman et al., 2021), leading to the accumulation of protein aggregates in lysosomes, defective mitochondria, and oxidative stress (Cuervo et al., 2005). In contrast, autophagy-stimulating interventions—such as caloric restriction, mTOR inhibition, or treatment with compounds like spermidine—extend lifespan and delay the onset of age-related phenotypes in diverse model organisms (Vellai et al., 2003; Harrison et al., 2009; Bitto et al., 2016; Alsaleh et al., 2020). These observations position autophagy as a key regulator of longevity (Aman et al., 2021; Pyo et al., 2013).

Immune system function is particularly dependent on autophagy (Deretic, 2006; Germic et al., 2019; Metur and Klionsky, 2021). With advancing age, immune function deteriorates, resulting in immunosenescence—a state characterized by reduced adaptive responses and chronic activation of innate immunity (Franceschi, 2000; Phadwal et al., 2012; Puleston and Simon, 2014). Defective autophagy in immune cells contributes to excessive cytokine release and metabolic dysfunction, reinforcing systemic inflammaging (Deretic, 2006; Aman et al., 2021). Similarly, in the central nervous system, the decline of autophagy impairs neuronal and glial homeostasis, leading to the accumulation of intracellular aggregates and activation of pro-inflammatory microglia (Boland et al., 2008; Caponio et al., 2022). Age-associated metabolic reprogramming shifts microglia toward a glycolytic, M1-like phenotype that sustains neuroinflammation and neuronal damage (Li et al., 2018). Enhancing autophagy across neural and immune compartments therefore represents a promising avenue to preserve cellular function and counteract age-related degeneration.

β-glucans are naturally occurring polysaccharides found in yeast, fungi, and cereals, long recognized for their immunomodulatory, antioxidant, and anti-inflammatory properties (Novak and Vetvicka, 2008; Kim et al., 2011; Murphy et al., 2020). β-glucans (BGs) act through the Dectin-1 receptor and related pathways to reprogram innate immune cells, improve pathogen defence and enhance tissue repair (Brown and Gordon, 2005). Recent evidence suggests that these molecules can also modulate mitochondrial respiration (Brogi et al., 2021) and autophagic flux (Li et al., 2019), linking immune regulation with metabolic resilience. However, their potential role in modulating the biology of aging, particularly within the brain, remains poorly characterized. Some positive actions of BGs on neural function in vivo were previously described, but it remains unclear whether BGs act on the CNS directly or the gut microbiome mediates their action.

The short-lived African turquoise killifish, *Nothobranchius furzeri*, provides a unique vertebrate model for experimental studies on aging. With a lifespan of only a few months, *N. furzeri* recapitulates many key cellular hallmarks of mammalian aging and in particular chronic activation of innate immunity, mitochondrial dysfunction, protein aggregation, oxidative stress, fibrosis, neuroinflammation and gliosis (Genade et al., 2005; Hartmann et al., 2009; Di Cicco et al., 2011; Van Houcke et al., 2021; Van Houcke et al., 2023). This model organism is therefore particularly convenient for studying the effects of dietary interventions on aging as it combines a lifespan typical of invertebrate models with the possibility of conducting fine-grained phenotypical investigations that include traits of direct translational relevance.

BGs have either a linear or a branched structure (Synytsya and Novak, 2013). Most studies investigated the action of linear β-glucans, linear BGs at high dosage were previously shown to exert positive effects in another *Nothobranchius* species. Here we investigated the effects of a realistic feed supplementation with the ramified 1,3–1,6 BGs on aging in vivo using N. furzeri. To investigate a direct action of BGs on the CNS, we exploited an ex vivo brain culture system complemented by investigations in human iPSC-derived cortical neurons. We hypothesized that BGs could mitigate aging-related dysfunction by enhancing autophagic and lysosomal activity, improving mitochondrial respiration, and reducing neuroinflammatory signalling. As components of the human diet, BGs are particularly relevant in the context of health promoting nutrition.

## RESULTS

### Effects of BGs in vivo

#### · Effects of diet fortified with BGs on aging phenotypes

*Nfu* were fed control diet (n=25) and diet fortified with 12.5 mg/kg (n=26) or 125 mg/kg (n=28) BGs, corresponding to the dose range commonly used in animal feed, from 2 to 27 weeks post-hatching when all surviving fish were sacrificed (n=10, n=9, and n=10, respectively). Survival analysis (Log-rank, χ2 = 0.2286) did not detect significant effects of BGs on mortality (Fig. 1A). Final body weight was significantly influenced by sex [p<0.0001, Two-way ANOVA; (Suppl. Fig. 1B)], but not treatment [p=0.4522, One-Way ANOVA; (Suppl. Fig. 1A)]. To define the effects of BGs on organ aging, we quantified age-associated phenotypes in brain, liver, muscle, heart and kidney for a total of 19 phenotypes (Table 1).

**Figure 1.**
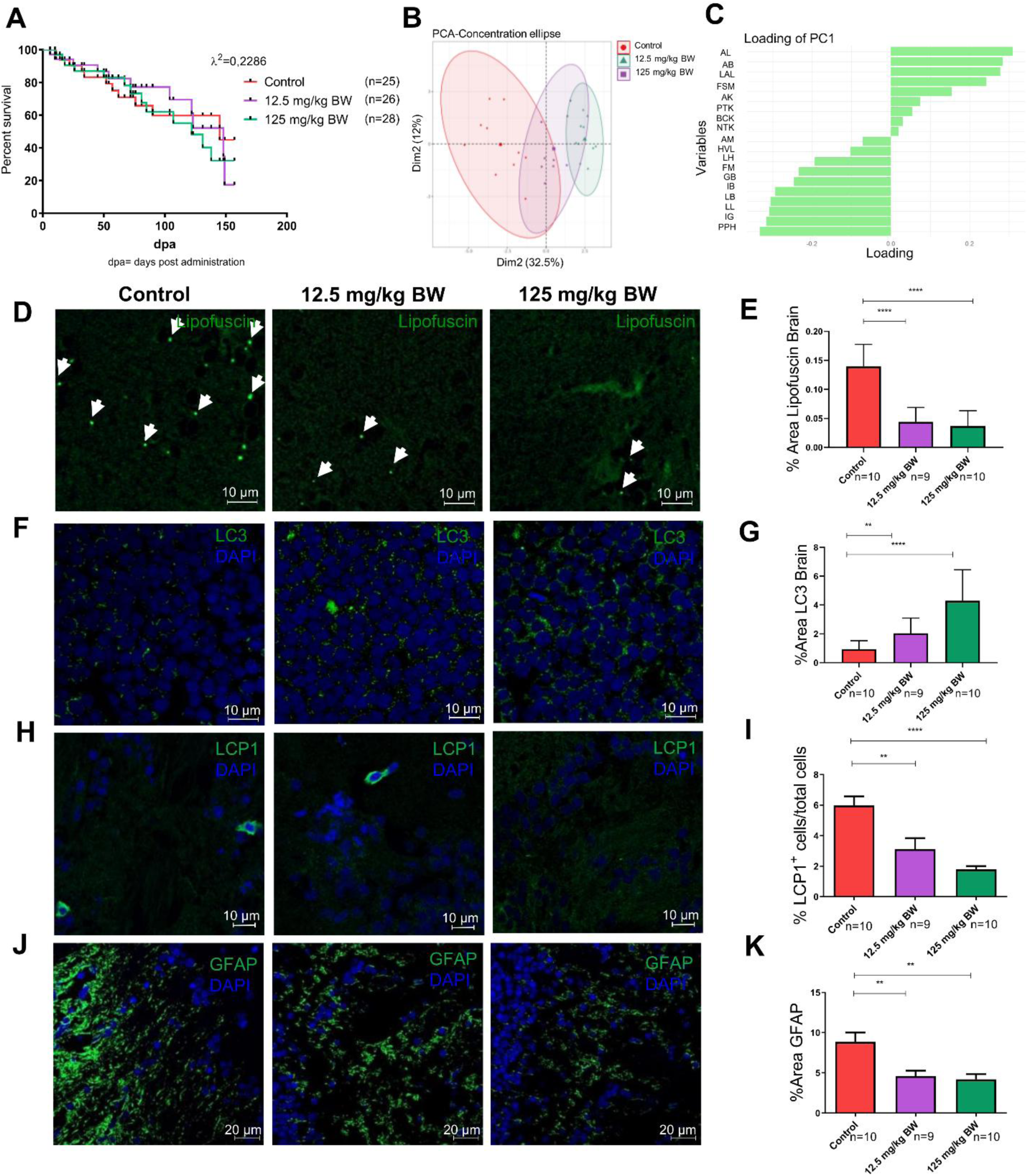
β-glucan supplementation improves survival and modulates aging hallmarks in the brain of *Nothobranchius furzeri*. (A) Kaplan–Meier survival curves of fish treated with β-glucans (BGs) at 12.5 mg/kg or 125 mg/kg body weight (BW) compared with control animals. Statistical significance was assessed using the log-rank (Mantel–Cox) test. (B) Principal component analysis (PCA) showing separation of samples based on BG treatment. Each point represents an individual sample. (C) Loading plot indicating the main variables contributing to principal component 1 (PC1). (D) Representative fluorescence images showing lipofuscin autofluorescence in brain sections from control, 12.5 mg/kg BW, and 125 mg/kg BW BG-treated fish. Arrowheads indicate lipofuscin granules. (E) Quantification of % area lipofuscin. (F) Representative confocal images of LC3 immunostaining (green) with DAPI nuclear staining (blue) in brain sections from control and BG-treated animals. (G) Quantification of % area LC3. (H) Representative images of LCP1 immunostaining (microglia marker) with DAPI counterstaining. (I) Quantification of % LCP1+ cells. (J) Representative images of GFAP immunostaining (astroglial marker) with DAPI staining. (K) Quantification of GFAP fluorescence intensity. Data are presented as mean ± SEM. Statistical significance was assessed using one-way ANOVA followed by Tukey’s multiple comparison test. Scale bars: 10 μm for Lipofuscin, LC3, LCP1 and 20 μm for GFAP. **Abbreviations:** LAL = Lysosomal activity Liver; PPH = Protein peroxidation Heart; AL = Autophagy Liver; LB = Lipofuscin Brain; IG = Inflammation Gut; PPL = Protein Peroxidation Liver; IB = Inflammation Brain; LL = Lipofuscin Liver; AB = Autophagy Brain; GB = Gliosis Brain; FM = Fibrosis Muscle; FSM = Fiber Size Muscle; LH = Lipofuscin Heart; AK = Area Kidney; PTK = Precipitated Tubules Kidney; NTK = Necrotic Tubules Kidney; HVL = Hepatocyte Vacuolization Liver; BCK = Bowman’s Capsule enlargement Kidney; AM = Autophagy Muscle. Control n = 10, 12.5 mg/Kg BW n = 9, 125 mg/Kg BW n = 10. *p < 0.05, **p < 0.01, ***p < 0.001.

**Table 1.** Aging phenotypes in different organs. Phenotypes, organs analyzed, p-values, FDR, and abbreviations used in the study.

| Aging Phenotype | Organ | pvalue | FDR | Abbreviation |
| --- | --- | --- | --- | --- |
| Lysosomal activity | Liver | 0.0001 | 0.0019 | LAL |
| Peroxidation protein | Heart | 0.0001 | 0.0009 | PPH |
| Autophagy | Liver | 0.0001 | 0.0006 | AL |
| Lipofuscin | Brain | 0.0001 | 0.0004 | LB |
| Inflammation | Gut | 0.0001 | 0.0003 | IG |
| Peroxidation protein | Liver | 0.0001 | 0.0003 | PPL |
| Inflammation | Brain | 0.0001 | 0.0002 | IB |
| Lipofuscin | Liver | 0.0003 | 0.0007 | LL |
| Autophagy | Brain | 0.0004 | 0.0008 | AB |
| Gliosis | Brain | 0.0008 | 0.0015 | GB |
| Fibrosis | Muscle | 0.0089 | 0.0153 | FM |
| Feret's size | Muscle | 0.0141 | 0.0223 | FSM |
| Lipofuscin | Heart | 0.0224 | 0.0327 | LH |
| Area renal tubules | Kidney | 0.0518 | 0.0703 | AK |
| Precipitation renal tubules | Kidney | 0.0519 | 0.0657 | PTK |
| Necrotic renal tubules | Kidney | 0.4116 | 0.4888 | NTK |
| Hepatocytes vacuolization | Liver | 0.5544 | 0.6196 | HVL |
| Bowman's capsule enlarged | Kidney | 0.7818 | 0.8252 | BCK |
| Autophagy | Muscle | 0.9395 | 0.9395 | AM |

Principal Component Analysis (PCA) was used to assess the global effect of BGs on these phenotypes (Fig. 1B). PC1 axis (32.5% variance) clearly separates control fish from those treated with 125 mg/kg Body Weight (BW) BGs, with the fish treated with 12.5 mg/kg BW BGs occupying an intermediate position, indicating a dose-dependent preventive effect of BGs on aging in *Nfu*. The loadings of PC1 (Fig. 1C) show that the variables with the greatest influence on group separation are Autophagy (Brain and Liver) and Lysosomal activity (Fig. 1C) among those increased by BGs and protein peroxidation (Heart and Liver), Inflammation (Brain and Gut), Lipofuscin (Liver and Brain), and Gliosis among those decreased by BG. BGs significantly modulated 12 phenotypes in the opposite direction compared to aging (FDR<0.05, Two-way ANOVA). Among these, lipofuscin in brain [p<0.0001, One-Way ANOVA; (Fig. 1D-E)], liver [p=0.0003, One-Way ANOVA; (Suppl. Fig. 1C-D)] and heart [p=0.0224, One-Way ANOVA; (Suppl. Fig. 1E-F)] in a dose-dependent manner and LC3 immunoreactivity in the brain [p=0.0004, One-Way ANOVA; (Fig. 1F-G)] and liver [p<0.0001, One-Way ANOVA; (Suppl. Fig. 1G-H)], suggesting the induction of autophagy. In the liver, BGs also significantly increased lysosomal activity (detected as colocalization of LAMP1 and Aggresome) in a dose-dependent manner [p<0.0001, One-Way ANOVA; (Suppl. Fig. 2A-B)], further supporting the modulation of autophagic processes and, consistently, lipofuscin accumulation was reduced in both brain and liver. However, no significant dose-dependent induction of autophagy by BGs was observed in muscle [p=0.9395, One-Way ANOVA; (Suppl. Fig. 1I-J)], although BGs were found to preserve muscle histology during aging. Indeed, 125 mg/kg BW of BGs significantly increased Feret’s size [p=0.0141, One-Way ANOVA; (Suppl. Fig. 3A-B)] while reducing fibrosis [p=0.0089, One-Way ANOVA; (Suppl. Fig. 3C-D)]. These results suggest a protective effect on muscle not mediated by autophagy induction. Furthermore, BG treatment at both concentrations significantly reduced the number of LCP1+ cells in the brain, indicating a reduction in neuroinflammation [p<0.0001, One-Way ANOVA; (Fig. 1H-I)]. A similar reduction in inflammation was observed in the gut [p<0.0001, One-Way ANOVA; (Suppl. Fig. 2C-D)], further supporting the anti-inflammatory effects of BGs across different tissues. Gliosis, a second marker of neuroinflammation, was significantly reduced in brain tissues of *Nfu* fed with BG-fortified feed in a dose-dependent manner [p=0.0008, One-Way ANOVA; (Fig. 1J-K)], providing independent support for an anti-inflammatory effect of BGs. In addition, peroxidation of proteins (Nitrotyrosine) was prominent in control livers (Suppl. Fig. 2E-F), and hearts (Suppl. Fig. 2G-H) and BGs significantly reduced this phenotype in both organs (p<0.0001 for both, One-Way ANOVA) in a dose-dependent manner. In contrast, BG treatment did not significantly modify hepatocyte vacuolization (steatosis) or the distribution of glycogen- and lipid-vacuoles in the liver (p=0.5544, One-Way ANOVA; Chi²=0.4475, p=0.7995; Supp. Fig. 3E-G). Similarly, BGs did not significantly affect age-related renal histopathological alterations, including Bowman’s capsule thickness (p=0.7818), incidence of precipitates in renal tubules (p=0.0519), necrotic renal tubules (p=0.4116) or tubular area (p=0.0518) (One-Way ANOVA; Supp. Fig. 4A-E). Our data indicate organ-specific BGs effects, with notable benefits observed in the brain, liver, heart and muscle but not in the kidneys.

#### Acute effects of diet fortified with BGs on brain aging

In order to test how rapidly BGs can exert their effects, we administered BGs at a concentration of 125 mg/kg BW for one week and analysed specifically the brain as this organ can be cultured *ex-vivo*, enabling mechanistic dissection of BGs action (see below). A total of 5 animals per group were used for both the treatment and control groups. BGs induced a statistically significant increase in LC3 immunoreactivity in the aged brain (padj=0.02475, t-test adjusted FDR), suggesting effective induction of autophagy during acute treatment, but failed to clear the lipofuscin accumulated over the course of the organism’s lifetime (padj=0.24216, t-test adjusted with False Discovery Rate or FDR), indicating that BGs can retard the accumulation of lipofuscin but fail to remove lipofuscin once it has accumulated (Fig. 2A and 2C). The magnitude of this LC3 induction, however, was not significantly different from that observed with chronic treatment (padj=0.09924, t-test adjusted with FDR), as shown in Figure 2B and 2D.

**Figure 2.**
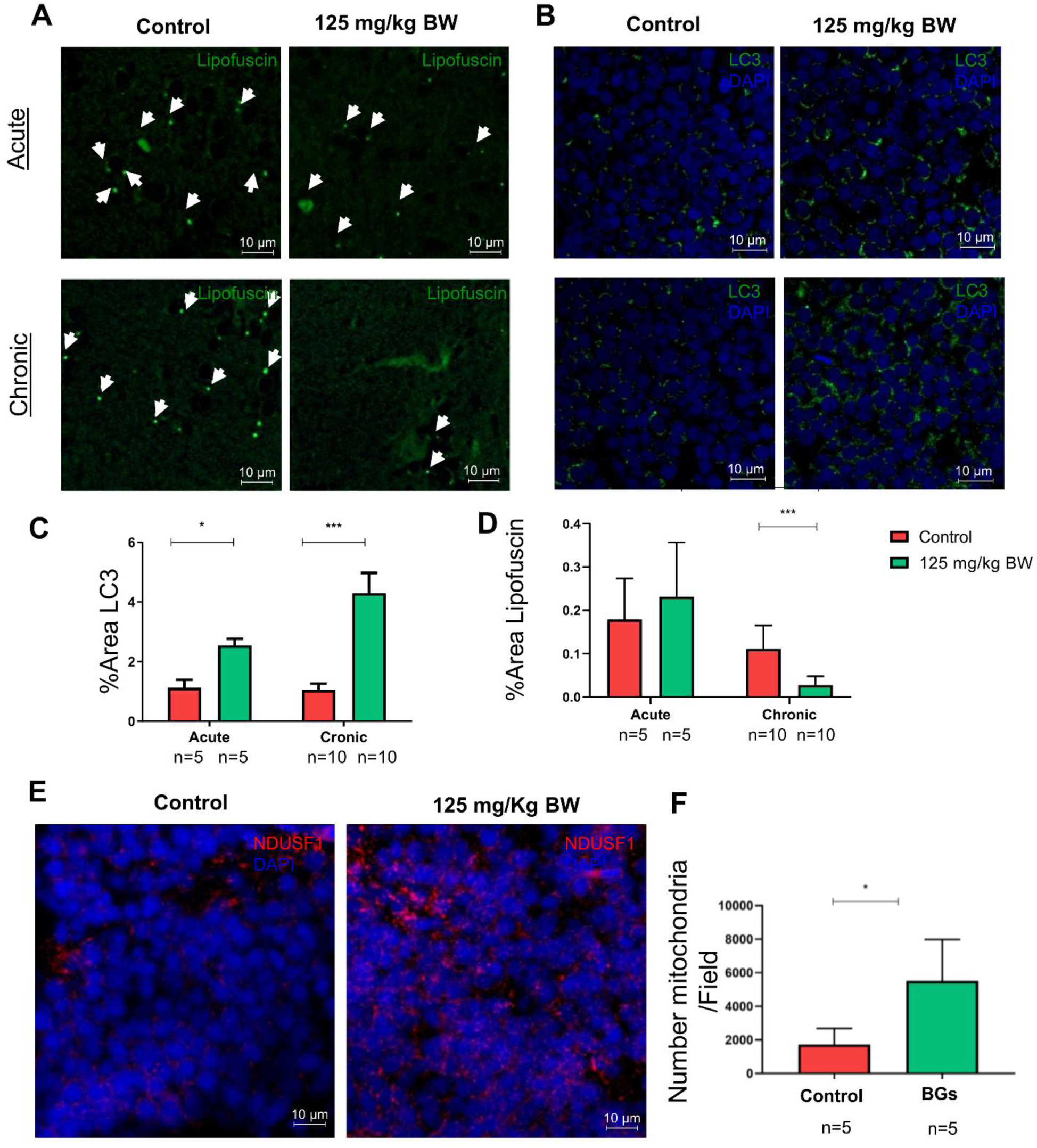
Acute BGs treatment modulates autophagy and mitochondrial markers in the brain of *Nothobranchius furzeri*. (A) Representative fluorescence images showing lipofuscin autofluorescence in brain sections from control and BG-treated fish (125 mg/kg BW) under acute and chronic treatment conditions. Arrowheads indicate lipofuscin granules. (B) Representative confocal images of LC3 immunostaining (green) with DAPI nuclear staining (blue) in control and BG-treated animals under acute and chronic conditions. (C) Quantification of % area of lipofuscin in acute and chronic treatment groups. (D) Quantification of % Area of LC3. (E) Representative confocal images of NDUFS1 immunostaining (red), a mitochondrial complex I marker, with DAPI counterstaining in control and BG-treated brains. (F) Quantification of NDUFS1 Area %. Data are presented as mean ± SEM. Statistical analysis was performed using unpaired two-tailed Student’s t-test with FDR correction for multiple comparisons (lipofuscin and LC3), unpaired two-tailed Student’s t-test for NDUFS1. Sample sizes: acute treatment n = 5 animals per group; chronic treatment n = 10 control, n = 9 (12.5 mg/kg BW), n = 9 (125 mg/kg BW). Scale bar: 10 μm. *p < 0.05, **p < 0.01, ***p < 0.001.

In addition, acute BG treatment significantly increased the number of mitochondria labelled with NDUFS1, a marker of mitochondrial complex I [p = 0.0129, t-test; (Fig. 2E–F)]. This result indicates that BGs promote mitochondrial biogenesis in the aged brain following short-term treatment.

### Effects of BGs on *ex-vivo* brains

#### BGs act directly on the brain and restore defective autophagy during aging

We investigated the direct effects of BGs on brain, we took advantage of our recently established protocol of killifish using *ex-vivo* brain cultures (Bagnoli et al., 2022). We cultivated half brains from fish of different ages (5, 10, 27 wph) and cultivated them for three days, exposing them at different combinations of compounds or vehicle. We concentrated our analysis on the optic tectum, because its periventricular gray zone is composed of tightly packed neuronal cells and is almost devoid of other cell types (Bagnoli et al., 2022). Aging led to a significantly affected LC3+ autophagosomes with reduction in the number [young vs. adult p=0.0003; young vs. aged p=0.0002; Two-way ANOVA; (Fig. 3A and 3C)], volume [young vs. adult p=0.0025, young vs. aged p=0.0010, Two-way ANOVA; (Fig. 3A and 3D)] and percentage area ([young vs. adult p=0.0318, young vs. aged p= 0.0027, Two-way ANOVA; (Fig. 3A and 3E)]. These findings suggest that autophagy is impaired in brain slices from adult and aged fish. No significant induction of LC3 immunoreactivity was observed in young slices treated with BGs [(p=0.9518, Two-way ANOVA; (Fig. 3A and 3C-E)]. On the contrary, BGs induced LC3+ autophagosomes in adult and aged brains, increasing the number [(young vs. adult p<0.0001; young vs. aged p<0.0001, Two-way ANOVA; (Fig. 3A and 3C)], volume [young vs. adult p=0.0092, young vs. aged p=0.0040, Two-way ANOVA; (Fig. 3A and 3D)], and percentage area of autophagosomes [young vs. adult p=0.0139; young vs. aged p=0.0018; Two-way ANOVA; (Fig. 3A and 3E)]. Thus, BGs can exert direct effects on the brain, normalizing autophagy impairments in adult and aged fish. Increased autophagosomes may result from two opposite processes: augmented autophagic flux and increased production of autophagosomes or an obstruction in the autophagic flow and accumulation of not processed autophagosomes. To confirm that BGs induce autophagy, we blocked autophagy with Bafilomycin A1 in *ex-vivo* brain slices from young brains (5 wph) inducing an accumulation of autophagosomes compared to the control [p<0.0001, One-way ANOVA; (Fig. 3B and 3F)] and BG treatment [p=0.001, One-way ANOVA; (Fig. 3B and 3F)], an increase in the percentage area of autophagosomes compared to the control [p=0.0159, One-way ANOVA; (Fig. 3B and 3G)] and BGs [p=0.0253, One-way ANOVA; (Fig. 3B and 3G)], and an increase in the volume of autophagosomes compared to the control [(p=0.0029, One-way ANOVA; (Fig. 3B and 3H)] and BGs [p=0.0023, One-way ANOVA; (Fig. 3B and 3H)]. The co-treatment with Bafilomycin A1 and BGs significantly augmented the number (p=0.0110, One-way ANOVA), volume (p<0.0001, One-way ANOVA), and percentage area (p = 0.0162, one-way ANOVA) of autophagosomes relative to Bafilomycin A1 alone (Fig. 3B and 3F-H). Experiments using the CYTO-ID kit in *ex-vivo* aged brain (27 wph) further validated these results (Suppl. Fig. 5A). The number [p=0.0344, t-test;(Suppl. Fig. 5A)], volume [p=0.0167, t-test; (Suppl. Fig. 5C)], and percentage area [p=0.0307, t-test; (Suppl. Fig. 5D)], of autophagosomes were all higher in combined BGs and Bafilomycin A1 treatments as compared with Bafilomycin A1 alone. These findings show that BGs enhance autophagosome formation in cultured brain slices.

**Figure 3.**
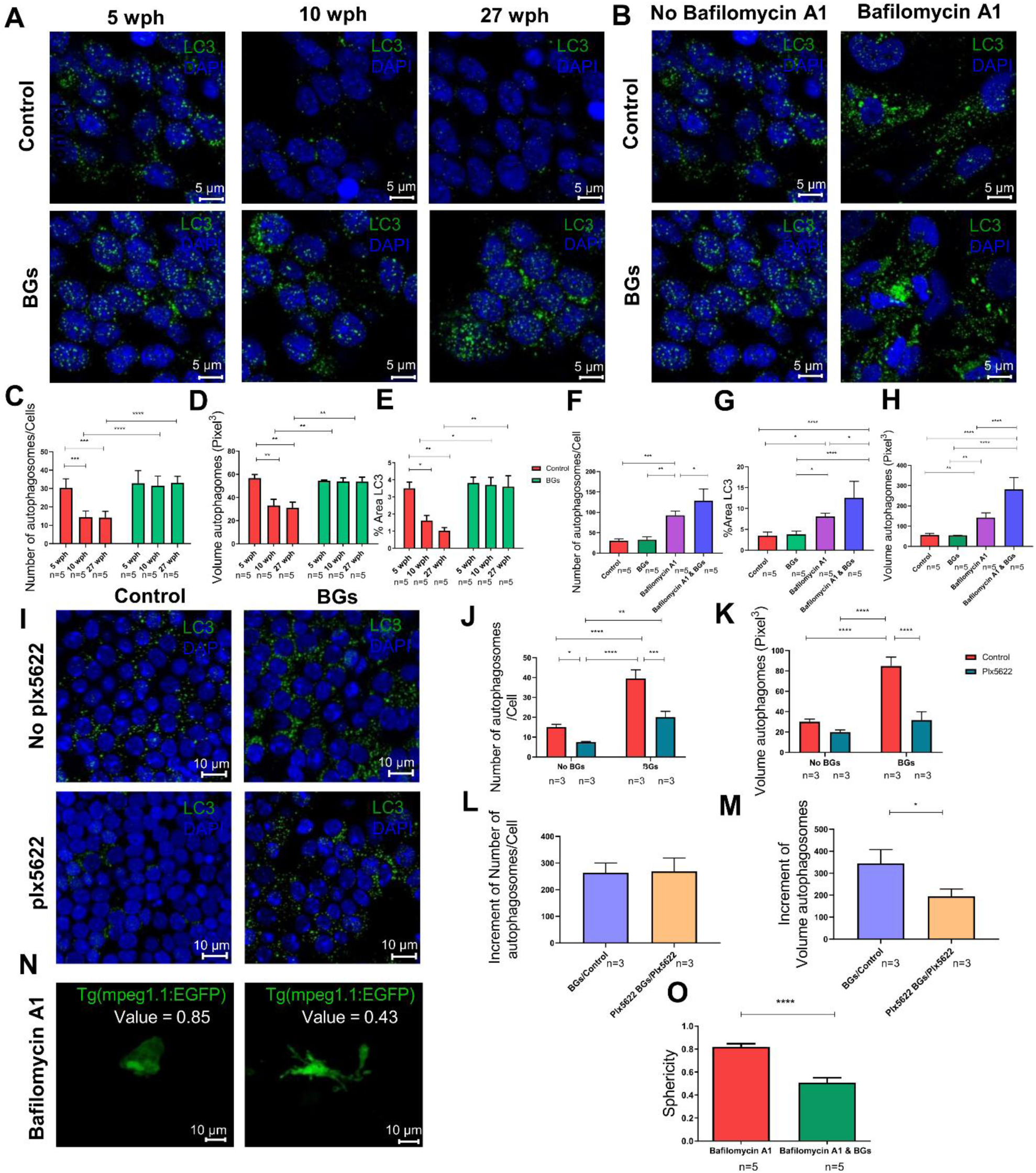
BGs restore impaired autophagy in the aging brain ex-vivo and partially act independently of microglia. (A) Representative confocal images of LC3 immunostaining (green) with DAPI nuclear staining (blue) in *ex-vivo* brain slices from *Nothobranchius furzeri* at 5, 10, and 27 weeks post hatching (wph) under control conditions or following β-glucan (BGs) treatment. Scale bar: 5 μm. (B) Representative confocal images of LC3 immunostaining in brain slices from 5 wph fish treated with Bafilomycin A1, with or without BGs treatment. Scale bar: 5 μm. (C) Quantification of autophagosome number per cell across age groups and treatment conditions. (D) Quantification of autophagosome volume across age groups and treatment conditions. (E) Quantification of percentage LC3-positive area across age groups and treatment conditions. n = 5 animals per group. (F) Quantification of autophagosome number per cell in control, BG-treated, Bafilomycin A1-treated, and BGs + Bafilomycin A1-treated explants. (G) Quantification of percentage LC3-positive area in control, BG-treated, Bafilomycin A1-treated, and BGs + Bafilomycin A1-treated explants. (H) Quantification of autophagosome volume in control, BG-treated, Bafilomycin A1-treated, and BGs + Bafilomycin A1-treated explants. n = 5 animals per group. (I) Representative confocal images of LC3 immunostaining in *ex-vivo* brain slices treated with plx5622 to deplete microglia, with or without BGs treatment. Scale bar: 10 μm. (J) Quantification of autophagosome number in control and microglia-depleted explants treated with BGs. (K) Quantification of autophagosome volume in control and microglia-depleted explants treated with BGs. n = 3 animals per group. (L) Quantification of the relative increase in autophagosome number induced by BGs in control and microglia-depleted slices. (M) Quantification of the relative increase in autophagosome volume induced by BGs in control and microglia-depleted slices. (N) Representative images of Tg(mpeg1.1: EGFP) zebrafish microglia treated with Bafilomycin A1 alone or in combination with BGs. Scale bar: 10 μm. (O) Quantification of microglial sphericity in Tg(mpeg1.1: EGFP) zebrafish following treatment with Bafilomycin A1 or Bafilomycin A1 + BGs. n = 5 animals. Data are presented as mean ± SEM. Statistical analyses were performed using unpaired two-tailed Student’s t-test, one-way ANOVA, or two-way ANOVA with Tukey’s post hoc test, as indicated. *p < 0.05, **p < 0.01, ***p < 0.001, ****p < 0.0001.

### Effects of BGs on global protein expression during aging

To obtain an unbiased molecular view of BGs effects, we employed proteomics and analyzed brain slices exposed to BGs to complement the targeted cellular analyses described above and provide a global picture of the molecular mechanisms underlying BGs effects.

Mass-spectrometry-based proteomics was employed to assess the effects of 3 days *ex-vivo* treatment with BGs on global protein regulation across three distinct age groups: 5-, 10-, and 27-wph. A total of approximately 4000 proteins were detected, with no significant differences observed between the three groups (χ² = 123; Fig. 4A). Similarly, the total number of differentially expressed proteins (DEPs) remained comparable across groups. However, a noticeable skew in favor of up-regulation emerged with advancing age (Fig. 4A). Notably, five proteins were significantly upregulated across all age groups: Aconitase 2 (ACO2), Myosin 10 (MYH10), Voltage-dependent anion-selective channel protein 1 (VDAC1), Pyruvate Carboxylase (PC), and Thioredoxin isoform 1 [TXT; (Fig. 4B)]. First, we examined whether BGs could restore dysfunctional pathways associated with aging *in vivo* by comparing our data with Di Fraia *et al*., (2025). A total of 337 proteins (∼9%) downregulated in aging were upregulated by BGs (Fig. 4C). Protein-Protein Interaction analysis [PPI; STRING, (PPI-enrichment p-value <7.52e^-8^, number edges = 1870, expected number edges = 1574)] identified four clusters enriched in Ribosome, Cytoskeleton and Synapses (Fig. 4D). Approximately 4.3% of proteins (58 proteins) that were upregulated during normal aging in vivo were downregulated by BGs. The PPI network showed a statistically significant enrichment but relatively sparse connectivity. Notably, a small cluster of proteins involved in metabolic pathways was observed, including ENO1, GAPDHS, FH, OGDH, UQCRC2, ASL, ALDH7A1 and CNDP1 [PPI-enrichment p-value < 8.05e−8; number of edges = 23; expected number of edges = 8; (Suppl. Fig. 6A)].

**Figure 4.**
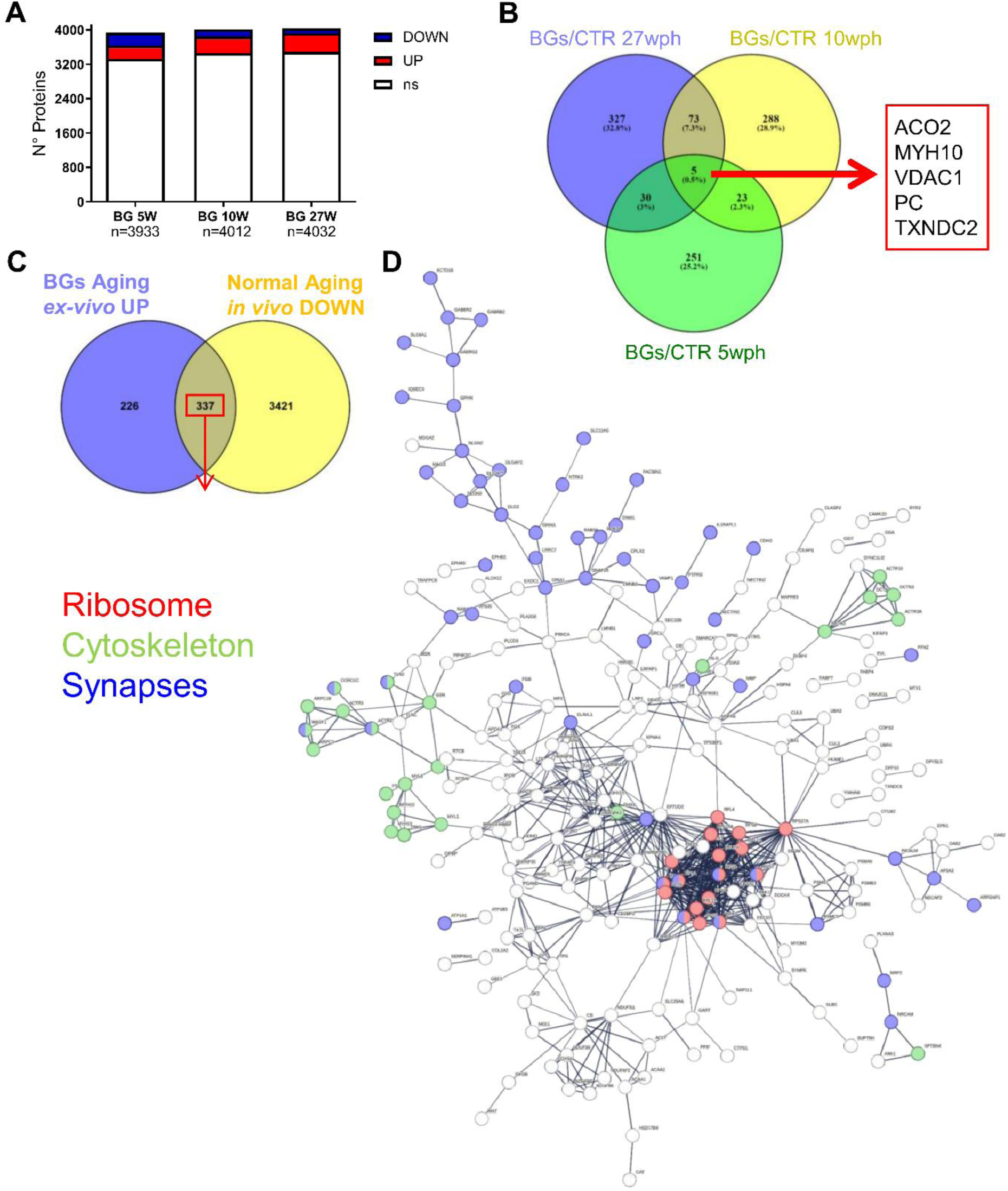
Proteomic analysis reveals pathways modulated by BGs that counteract molecular signatures of brain aging. (A) Distribution of differentially expressed proteins identified by proteomic analysis comparing 5, 10, 27-week post hatching (wph) after BGs treatment in *ex-vivo*. (B) Venn diagram showing the overlap of proteins significantly regulated by BG treatment relative to controls at 5, 10, and 27 (wph). Five proteins were commonly regulated across all comparisons. (C) Comparison between proteins upregulated by BG treatment *ex-vivo* in old brain and proteins downregulated during normal aging in vivo, revealing an overlap of 337 proteins, suggesting that BG treatment partially reverses age-associated molecular changes. (D) Protein–protein interaction network analysis of the overlapping protein set. Functional clustering highlights pathways related to ribosome function (red), cytoskeletal organization (green), and synaptic processes (blue). Network visualization was generated using protein–protein interaction databases and pathway enrichment analysis, illustrating functional modules affected by BG treatment in the aging brain. wph = weeks post hatching.

Additionally, we investigated the effects of BGs on aged brain (at 27 wph) *ex-vivo*. BGs upregulated proteins associated with Mitochondrial respiration, particularly complex I, Synapses, Axons and Cytoskeleton, as highlighted by PPI [PPI-enrichment p-value <1e^-16^, number edges = 4869, expected number edges = 4164; (Fig. 5A)]. In particular, increased expression of proteins associated with lysosomal lumen acidification (ATP6V0A1, ATP6V0D1, ATP6V0E1, ATP6V1H, ATP6V1F) was detected (Figure 5A). Finally, the proteins downregulated by BGs were fewer than those upregulated. Due to the limited number of downregulated proteins, no statistically significant (PPI-enrichment p-value = 0.138, number edges = 68, expected number edges = 59) PPI enrichments were detected (Suppl. Fig. 6B).

**Figure 5.**
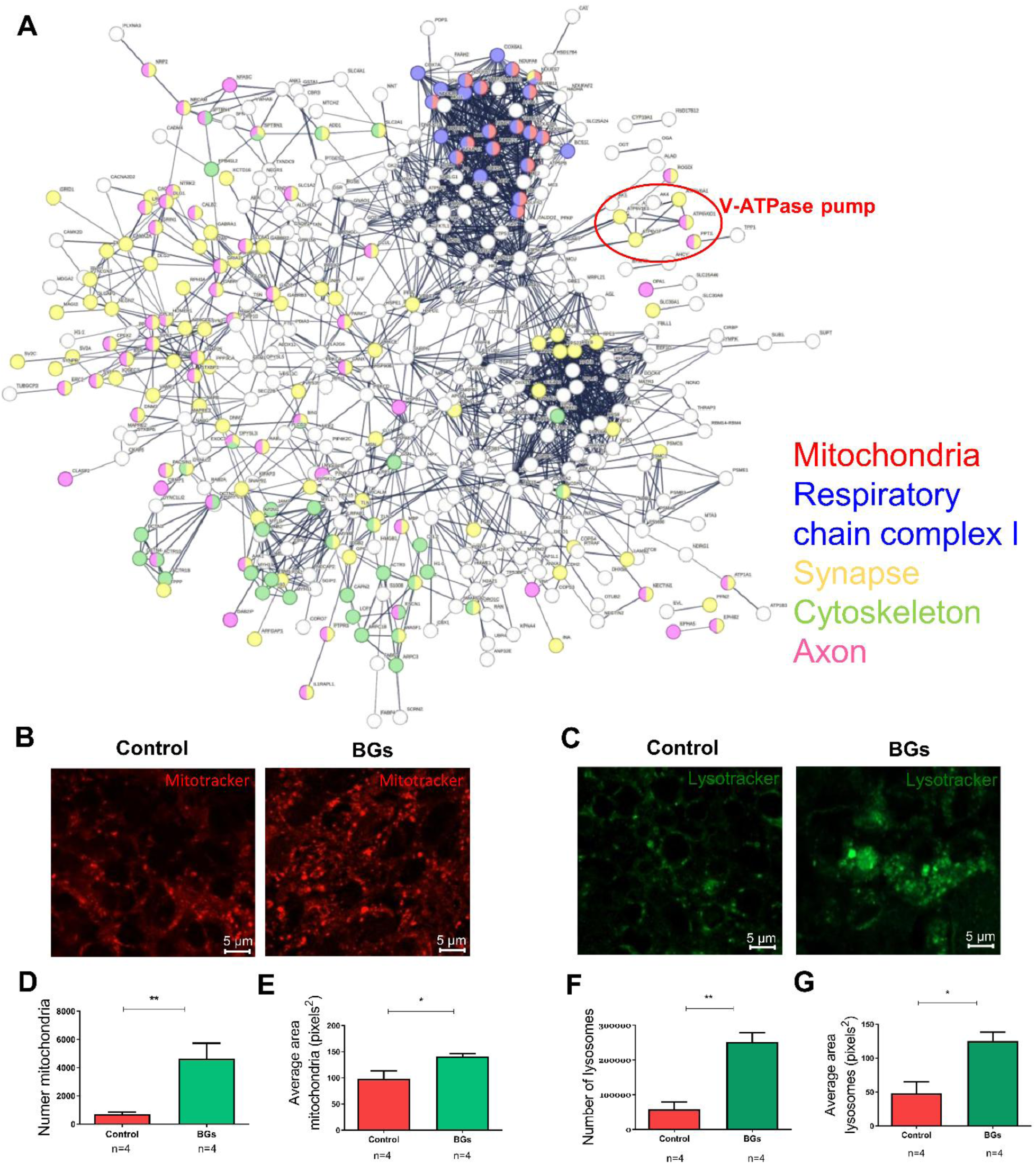
BGs treatment at 27 wph promotes mitochondrial and lysosomal pathways in the aging brain. (A) Protein–protein interaction network of proteins upregulated by BGs treatment at 27 wph identified by proteomic analysis. Functional clustering highlights pathways related to mitochondria (red), respiratory chain complex I (blue), synapse (yellow), cytoskeleton (green), and axon (magenta). A distinct cluster corresponding to the V-ATPase proton pump complex, involved in lysosomal acidification, is indicated. (B) Representative confocal images of MitoTracker staining showing mitochondrial activity in brain explants from control and BG-treated animals at 27 wph. Scale bar: 5 μm. (C) Representative confocal images of Lysotracker staining indicating lysosomal activity in control and BG-treated brain explants at 27 wph. (D) Quantification of Number Mitochondria. (E) Quantification of average area Mitochondria. (F) Quantification of lysosomes number. (G) Quantification of average area lysosomes. Data are presented as mean ± SEM. Statistical significance was determined using unpaired two-tailed Student’s t-test. Scale bar: 5 μm. *p < 0.05, **p < 0.01.

BGs targeted Oxidative phosphorylation, Lysosomal, and GABAergic synapse pathways, consistent with their protective effects against aging *ex-*vivo (Suppl.Fig. 7A–C).

At 10 wph, BG treatment upregulated proteins associated with Synapses and Axons terms (PPI-enrichment p-value <1e^-16^, number edges = 2796, expected number edges = 2131), in Suppl. Fig. 8A. Although not reaching statistical significance due to the limited number of proteins detected, a consistent trend toward upregulation of V-ATPase subunits (ATP6V0A1, ATP6V0D1, ATP6V0E1, ATP6V1H, ATP6V1F) was observed, suggesting a possible increase in lysosomal acidification activity. Notably, four of these subunits (ATP6V0A1, ATP6V0D1, ATP6V0E1, and ATP6V1H) were also found to be regulated by BGs at 27 wph, supporting a conserved modulation of lysosomal machinery across aging stages.

At 5 wph, when autophagy is not yet impaired, BG treatment did not induce significant enrichment of important intracellular pathways (PPI-enrichment p-value <1.1 e^-10^, number edges = 625, expected number edges = 479). Importantly, none of the proteins associated with lumen acidification or V-ATPase activity were found to be modulated by BGs at 5 wph, further supporting the view that the effects of BGs on lysosomal regulation emerge at later aging stages.

In order to validate the proteomic analysis, which showed that BGs induced proteins involved in mitochondrial respiration (Fig. 5D-E) in *ex-vivo* aged brain, we examined the morphology and number of mitochondria by MitoTracker staining. BG treatment significantly increased both the number [p=0.0071, t-test; (Fig. 5D)] and area [p=0.0366, t-test; (Fig. 5E)] of mitochondria in *ex-vivo* aged brain as compared to the controls. To complement the autophagy induction analysis, we focused on the morphology and number of acidic lysosomes (Fig. 5C and 5F-G) in ex- vivo aged brain (27 wph). BG treatment significantly increased both the number [p=0.0014, t-test; (Fig. 5F)] and area [p=0.0113, t-test; (Fig. 5G)] of acidic lysosomes visualized with Lysotracker.

#### BGs act directly on microglia and neurons

We sought to determine the identity of the cells responding to BGs, focusing on microglia and neurons.

### • BGs reduce microglia activation

To analyze effects of BGs reduce microglial activation, we prepared *ex-vivo* brain slices from young adult zebrafish Tg(mpeg1.1:EGFP) enabling precise analysis of microglial morphology. To induce dysfunctional autophagy and activate the microglia, we treated the slides with Bafilomycin A1. BGs significantly reduced the sphericity [p<0.0001, t-test; (Fig. 3N-O)] of microglial cells in Tg(mpeg1.1:EGFP) zebrafish after Bafilomycin A1 treatment (Fig. 3N-O).

### • Induction of neuronal autophagy by BGs is partially mediated by microglia

The previous result aligns with a substantial body of evidence indicating that BGs act on immune cells, particularly microglia and macrophages (Shah et al., 2008; Goodridge et al., 2009; De Marco Castro et al., 2020; Heng et al., 2021). To investigate whether BGs can affect neurons directly, we treated *ex-vivo* brain slices from *Nfu* with plx5622, a drug that depletes microglia (Stonedahl et al., 2022; Liu et al., 2019; Spangenberg et al., 2019). The percentage of LCP1^+^/total cells was significantly reduced [p=0.007, t-test; (Suppl. Fig. 5F)] in brain slices treated with 20 μM plx5622 as compared to controls (Suppl. Fig. 5E-F). We analysed the effect of BGs on autophagy in microglia-depleted *ex-vivo* brain slices where autophagy is dysregulated (Fig. 3I and 3J-M). Consistent with our previous experiments, treatment of adult brain slices with BG for 3 days increased the number of autophagosomes per cell [p<0.0001, Two-way ANOVA; (Fig. 3J)]. Microglial depletion reduced the number of autophagosomes per cell in control slices [p=0.0385, Two-way ANOVA; (Fig. 3J)], but did not alter autophagosome volume [p=0.2615, Two-way ANOVA; (Fig. 3K)]. BGs increased the number of autophagosomes in microglia-depleted slices [p=0.0022, Two-way ANOVA; (Fig. 3J)], but not to the level observed in control slices treated with BGs [p=0.0001, Two-way ANOVA; (Fig. 3J)] and did not induce a significant increase in autophagosome volume [p=0.1774, Two-way ANOVA; (Fig. 3K)]. When the baseline condition is considered, the relative increase in the number of autophagosomes did not differ significantly between control and microglia-depleted slices [p=0.8984, Two-way ANOVA; (Fig. 3L)]. In contrast, the relative increase in autophagosome volume induced by BGs was significantly larger in control slices than in microglia-depleted slices [(p=0.0215, t-test; (Fig. 3M)]. These results suggest that the effects of BGs on neurons were only partially mediated by microglia.

### BGs improve survival of human *iPSC*-derived *neurons after autophagy inhibition*

Since previous results have shown that the effects of BGs in *Nfu* were not entirely mediated by microglia, we studied the effect of BGs on human iPSC-derived cortical neurons. Considering that these human iPSC-derived cortical neurons (90 DIV) are equivalent to embryonic neurons and do not suffer from autophagy impairment, we treated them with Bafilomycin A1 to induce autophagy impairment.

Treatment with Bafilomycin A1 (10 nM and 100 nM) for 72 hours significantly reduced the number of viable neurons as compared to control [p < 0.0001, two-way ANOVA; (Fig. 6A-B)]. However, treatment with 100 nM Bafilomycin A1 for 24 hours did not induce significant neuronal mortality as compared to the control [p = 0.5189, two-way ANOVA; (Suppl. Fig. 9A-B)]. BGs significantly reduced neuronal mortality in neurons treated for 72 hours with 10 nM Bafilomycin A1 ([p=0.037, Two-way ANOVA; (Fig. 6A-B)], but not with 100 nM Bafilomycin A1 [p=0.3644, Two-way ANOVA; (Fig. 6A-B). Moreover, BGs did not improve neuronal survival under conditions where mortality was not elevated, such as in the control condition [p=0.7417, Two-way ANOVA; (Fig. 6A-B)] or during treatment with 100 nM Bafilomycin A1 for 24 hours [p=0.9573, Two-way ANOVA; (Suppl. Fig.9A-B)].

**Figure 6.**
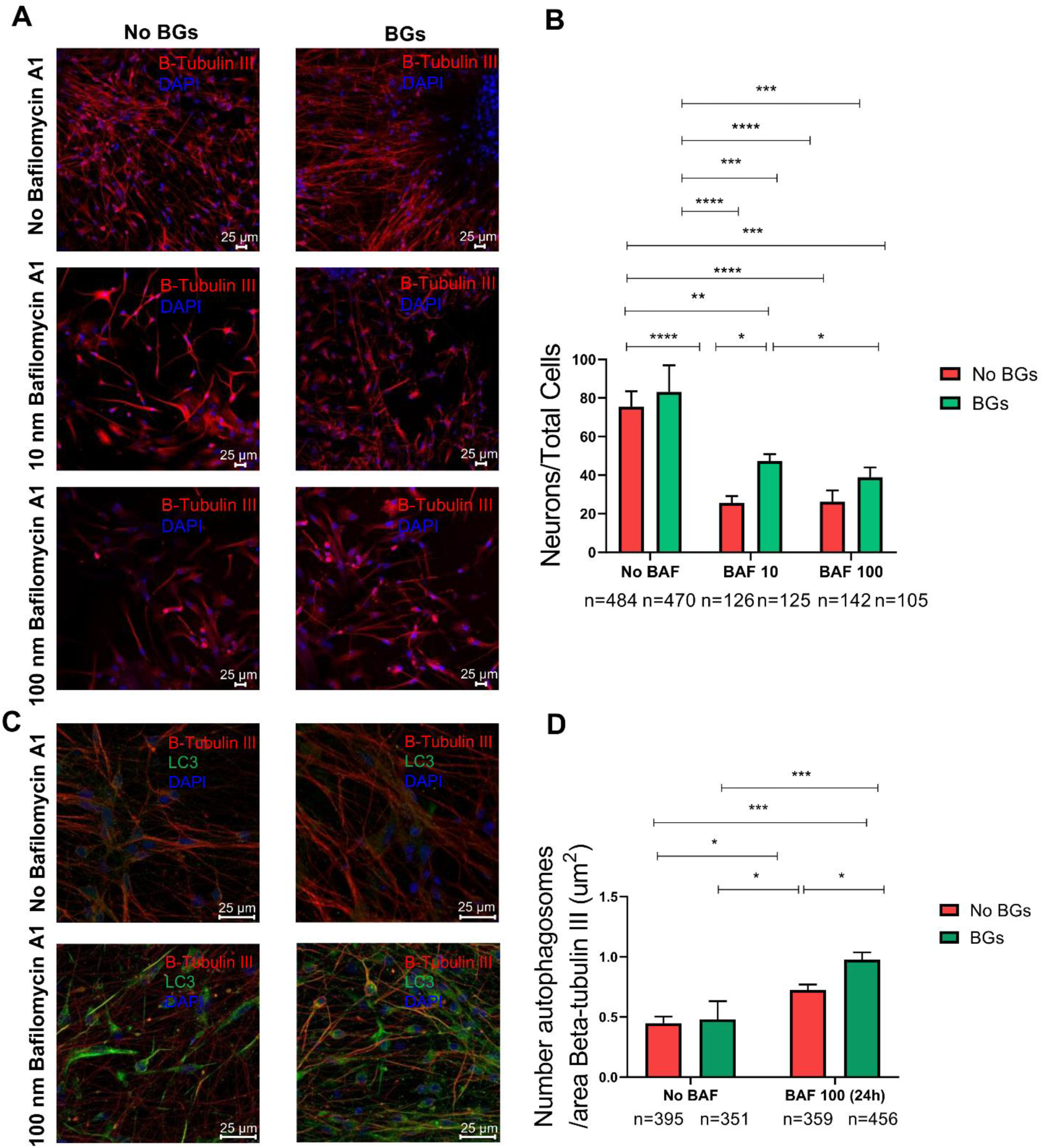
β-glucans improve survival and restore autophagy in human iPSC-derived cortical neurons under autophagy impairment. (A) Representative confocal images of human iPSC-derived cortical neurons stained for β-Tubulin III (red) and DAPI (blue) under control conditions, 10 nM Bafilomycin A1 treatment, and 100 nM Bafilomycin A1 treatment, with or without β-glucans (BGs). Scale bar: 25 μm. (B) Quantification of neuronal survival expressed as Neurons/Total Cells ratio in control, Bafilomycin A1-treated, and BG-treated conditions. (C) Representative confocal images showing LC3 immunostaining (green), β-Tubulin III (red), and DAPI (blue) in neurons treated with 100 nM Bafilomycin A1 for 24 h, with or without BGs treatment. Scale bar: 25 μm. (D) Quantification of LC3-positive autophagosomes normalized to β-Tubulin III area (μm²). Data are presented as mean ± SEM. Statistical analysis was performed using two-way ANOVA followed by Tukey’s multiple comparison test. *p < 0.05, **p < 0.01, ***p < 0.001, ****p < 0.0001.

### BGs counteract impaired autophagy in *iPSC-*derived *neurons*

BGs did not induce autophagy under normal conditions where neurons were not perturbed [p=0.9918, Two-way ANOVA; (Fig. 6C-D)], consistent with the observation in *Nfu*, where BGs only exert an effect when autophagy is compromised. Both doses of Bafilomycin A1 led to a reduction in LC3 compared to control [p=0.0102; p=0.0482 for increasing doses, respectively, Two-way ANOVA; (Suppl. Fig. 10A-B)]. In contrast, 24 hours treatment with 100 nM Bafilomycin A1 increased the LC3/β-tubulin area ratio compared to untreated control neurons [p=0.0210, Two-way ANOVA; (Fig. 6C-D)]. The combined treatment of Bafilomycin A1 and BGs increased the LC3/β-tubulin area ratio compared to Bafilomycin A1 alone [p=0.0343, Two-way ANOVA; (Fig. 6C-D)]. These results indicate that BGs enhance autophagosome formation in human neurons under conditions where autophagy is compromised, confirming the findings in *Nfu* slices. This also suggests that BGs can exert a direct effect on human neurons.

## DISCUSSION

Our study investigated the effects of β-glucans (BGs) on multiple aging hallmarks using a combination of three experimental systems: in vivo dietary supplementation in killifish, *ex-vivo* organotypic cultures of killifish brain, and iPSC-derived human neurons. Using this combination of models, we observed a broad beneficial effect on aging hallmarks in peripheral organs as well as reduced neuroinflammation, increased neuronal autophagy–lysosome function, and induction of mitochondrial biogenesis. Notably, even though microglia show a response to BGs, the effects are partially mediated by a direct action of BGs on both killifish and human neurons.

A key finding of this work is the induction of autophagy and the attenuation of age-associated accumulation in cardiomyocytes, hepatocytes, and neurons of oxidized proteins and/or lipofuscin, a largely undegradable by-product of oxidative damage that progressively compromises lysosomal efficiency and autophagic clearance (Terman and Brunk, 2004; Cuervo et al., 2005; Kaushik and Cuervo, 2015). This result is consistent with the concept, established in mammalian systems, that maintenance of lysosomal function is a limiting factor for preserving proteostasis and cellular viability during aging (Cuervo et al., 2005; Nixon and Yang, 2012). Importantly, our data further suggest that the timing and duration of intervention is critical: while acute BGs treatment induced autophagy, only chronic supplementation dampened lipofuscin accumulation, supporting prior observations that late-life induction of autophagy may be insufficient to reverse the long-term buildup of lysosome-resistant material (Terman and Brunk, 2004; Kaushik and Cuervo, 2015). This aligns with studies showing that sustained restoration of lysosomal competence—rather than transient autophagy initiation—is required to achieve durable improvements in cellular quality control.

Importantly, accumulating experimental evidence indicates that physiological brain aging is accompanied by a progressive decline in lysosomal acidification, resulting in elevated lysosomal pH, reduced hydrolase activity, and impaired degradation even in the absence of overt neurodegenerative pathology (Nixon and Yang, 2012; Colacurcio and Nixon, 2016; Burrinha et al., 2023). Age-dependent reductions in V-ATPase assembly and proton-pumping efficiency have been proposed as central mechanisms underlying this deacidification, thereby establishing lysosomal pH dysregulation as a primary driver of proteostatic stress rather than a secondary consequence (Settembre et al., 2013; Wong et al., 2020). In line with this notion, is the observation that killifish brain aging is associated with loss of V-ATPase complex stochiometry with likely functional consequences on lysosomal acidification (Kelmer Sacramento et al., 2020). Notably, experimental re-acidification of aged neuronal lysosomes restores proteolytic competence and rescues synaptic integrity, underscoring lysosomal acidity as a functionally limiting determinant of neuronal resilience (Burrinha et al., 2023). Within this framework, the BG-dependent enhancement of Lysotracker staining (a dye sensitive to lysosomal acidification) observed in our models suggests that β-glucans act at a critical late-stage node of the autophagy–lysosome pathway. This notion is further supported by the upregulation of V-ATPase subunits. Rather than solely promoting autophagosome formation, BGs appear to restore degradative capacity by relieving pH-dependent constraints on cargo breakdown, thereby enabling efficient completion of autophagic and mitophagic flux. Given that impaired acidification has been causally linked to mitochondrial dysfunction, ROS accumulation, and inflammatory activation in aging systems (Nixon and Yang 2012; Kaushik and Cuervo, 2015; Yambire et al., 2019), restoration of lysosomal pH homeostasis may represent a unifying mechanism underlying the multi-organ protective effects observed in this study.

The *ex-vivo* organotypic brain slice model further allowed us to compare BG effects across ages in a brain-autonomous context. Similar to observations in mammalian tissues, aged slices exhibited reduced autophagic flux and altered mitochondrial morphology, consistent with the tight coupling between autophagy, mitophagy, and mitochondrial respiration reported across species (Green et al., 2011; Sun et al., 2016; Wilson et al., 2023). Importantly, autophagy is not merely a degradative pathway but directly regulates cellular bioenergetics by controlling the turnover of metabolic enzymes, mitochondrial components, and nutrient reservoirs required to sustain oxidative phosphorylation. Genetic or pharmacological inhibition of autophagy impairs mitochondrial respiration, reduces ATP production, and alters substrate utilization, underscoring that autophagic flux is metabolically permissive in addition to cytoprotective (Green et al., 2011; Sun et al., 2016; Wilson et al., 2023). Recent work has further demonstrated that the autophagy–NAD axis represents a critical interface between lysosomal recycling and mitochondrial redox metabolism, whereby impaired autophagy leads to NAD depletion, reduced respiratory capacity, and energetic decline in aging systems (Wilson et al., 2023). In aging tissues, defective autophagy therefore generates a dual energetic deficit: accumulation of dysfunctional mitochondria and reduced recycling of metabolic intermediates necessary for efficient respiration.

Acute in vivo BG administration was sufficient to induce early improvements in mitochondrial signatures, mirroring the rapid mitochondrial responses observed in the *ex-vivo* brain slice model and suggesting that mitochondrial homeostasis represents an early and sensitive target of BG-mediated intervention. Considering that lysosomal proton pumping is an ATP-dependent process and tightly coupled to mitochondrial respiration and NAD availability, maintenance of bioenergetic integrity appears essential for preservation of lysosomal acidification during aging. Our data showing BG-dependent improvement of mitochondrial and lysosomal morphology are consistent with studies demonstrating that restoration of autophagic competence indirectly normalizes mitochondrial function by promoting removal of damaged organelles, preserving NAD redox balance, and reducing ROS-driven inflammatory signaling (Sun et al., 2016; Aman et al., 2021; Wilson et al., 2023). Proteomic modulation of lysosomal acidification machinery, including V-ATPase-related components, further strengthens this mechanistic interpretation, as V-ATPase integrity and lysosomal acidification are essential for autophagosome maturation and efficient cargo degradation (Settembre et al., 2013; Zhang et al., 2016; Wong et al., 2020). This is consistent with the broader literature supporting lysosomal–autophagic competence as a key determinant of neuronal resilience and aging trajectories (Nixon and Yang, 2012; Kaushik and Cuervo, 2015).

Importantly, our proteomic analysis identified five proteins consistently upregulated across three different ages following BG treatment—ACO2, MYH10, VDAC1, PC, and TXNDC2—highlighting the engagement of a conserved mitochondrial quality-control network. ACO2 and PC sustain tricarboxylic acid cycle flux and metabolic flexibility, thereby supporting mitochondrial respiration, NAD redox homeostasis, and ATP production required for energy-dependent processes such as lysosomal proton pumping (Mangialasche et al., 2015; Cappel et al., 2019; Chen et al., 2020; Gondáš et al., 2023; Zhu et al., 2023; Padalko et al., 2024; Fukawa et al., 2025). VDAC1 regulates mitochondrial metabolite exchange and participates in PINK1/Parkin-dependent mitophagy signalling, directly linking mitochondrial status to autophagic clearance (Geisler et al., 2010; Sun et al., 2012; Ham et al., 2020; Argueti-Ostrovsky et al., 2024). MYH10 contributes to cytoskeletal dynamics required for autophagosome trafficking and lysosomal positioning, processes essential for efficient autophagosome–lysosome fusion (Javier-Torrent et al., 2020; Jun et al., 2023). TXNDC2, a thioredoxin family member, contributes to redox buffering and protection against oxidative stress, a critical determinant of both mitochondrial integrity and lysosomal membrane stability (Wang et al., 2016; Cha et al., 2023). Given emerging evidence that autophagy-dependent recycling sustains NAD availability and mitochondrial respiratory capacity during aging (Wilson et al., 2023), the coordinated regulation of these proteins supports a model in which BGs reinforce mitochondrial metabolism, redox homeostasis, and organelle turnover in an integrated manner. Given that mitophagy completion strictly depends on fully acidified lysosomes for degradation of mitochondrial cargo (Settembre et al., 2013; Wong et al., 2020), the parallel enhancement of lysosomal acidification and mitochondrial-associated proteomic signatures strongly suggests that BGs restore the autophagy–lysosome–mitochondria axis at multiple hierarchical levels. This systems-level reinforcement may explain the observed attenuation of oxidative stress and neuroinflammatory activation.

A major strength of our approach is the integration of neuroinflammatory readouts with autophagy modulation. Aging is associated with a progressive shift of microglia toward pro-inflammatory states that contribute to synaptic dysfunction and impaired tissue repair (Gomez-Nicola and Perry, 2015; Hammond et al., 2019; Sierra et al., 2019). Consistent with this literature, we observed microglial activation in aged brains and its attenuation upon β-glucan treatment. plx5622-based microglial manipulation allowed direct comparison with established CSF1R-inhibition studies in mammalian models and demonstrated that microglia contribute to—but do not fully account for—the enhancement of neuronal autophagy, consistent with a model in which BGs act through both immune-mediated and cell-autonomous mechanisms.

The translational relevance of these findings is anticipated by data obtained in human iPSC-derived cortical neuron. In line with extensive evidence linking lysosomal dysfunction and impaired autophagy to Alzheimer’s and Parkinson’s disease pathogenesis (Nixon and Yang, 2012; Menzies et al., 2015; Lou et al., 2019), BGs enhanced neuronal resilience under autophagy inhibition and partially restored autophagic activity following lysosomal blockade. These data place BGs within a broader landscape of interventions aimed at boosting proteostasis and lysosomal competence, including strategies targeting TFEB-dependent lysosomal biogenesis and autophagic flux (Settembre et al., 2013; Hansen et al., 2018).

Finally, the use of the short-lived vertebrate *Nothobranchius furzeri* provides a comparative perspective that complements mammalian aging studies (Terzibasi Tozzini et al., 2007; Cellerino et al., 2016). Our findings support the notion that interventions targeting conserved maintenance pathways—particularly the lysosomal acidification–autophagy–mitochondria axis—can produce beneficial effects across species. While future studies will be required to define tissue-specific responses and long-term outcomes in mammalian systems, our data collectively support β-glucan supplementation as a physiologically relevant approach to promote healthy aging and preserve brain function by restoring lysosomal acidification, reinforcing mitochondrial quality control, and attenuating inflammaging.

## ACKNOWLEDGEMENTS

The authors gratefully acknowledge Dr. Cinzia Caterino (Leibniz Institute on Aging – Fritz Lipmann Institute, Jena) for technical support. We thank the Proteomics Facility at the Leibniz Institute on Aging – Fritz Lipmann Institute for proteomic analyses, with special thanks to Dr. Emilio Cirilli for his support. The Tg(mpeg1.1) transgenic line was kindly provided by Prof. Ambrosio (Biogem, Ariano Irpino, Italy).

## MATERIAL AND METHODS

### Ethics statement

All experimental procedures were conducted in accordance with European Union Directive 2010/63/EU for the protection of animals used for scientific purposes and were approved by the Institutional Animal Care and Use Committee (IACUC) of the University of Pisa (authorizations B290E.N.VIC and B290E.N.TUN). Experiments were performed at Bio@SNS (Scuola Normale Superiore, Pisa), AquaLab (Department of Veterinary Sciences, University of Pisa) and Zebralab (IRCCS Stella Maris). Allocation of animals to experimental groups was randomized when applicable, and investigators performing image acquisition and quantitative analyses were blinded to experimental conditions.

### Nothobranchius furzeri Care and Maintenance

All animal experiments were performed on *Nothobranchius furzeri (Nfu)* strain MZCS-NF222 (Blažek et al., 2017). Embryos were hatched and housed locally in a recirculating Tecniplast Zebtech system with automatized water flow, pH and salinity control. Water parameters were set temperature of 28 °C, pH 7 and conductivity 800 μs/cm. The photoperiod was set to 12 h light 12 h dark in phase with external day/night cycle (Nath et al., 2023; Polacik et al., 2016; Dodzian et al., 2018). Fertilized eggs were maintained on wet peat moss at a temperature of 28 °C in sealed Petri dishes. When embryos reached the final stage of development, eggs were hatched by transferring them in 1: 1 humic acid (1g /L) at temperatures of 4 °C. Hatched embryos were transferred to a clean vessel with static water with daily water changes where they were kept until 2 weeks of age. For the first two weeks, fry were fed with newly hatched *Artemia nauplii* and were maintained in the incubator with the same parameters mentioned above. After age 2 weeks the diet was gradually shifted to frozen *Chironomus* larvae. In vivo experimental design.

### Zebrafish Care and Maintenance

The study was carried out using Tg(mpeg1.1: EGFP) zebrafish (Ellett et al., 2011; Renawat and Masai, 2021; Ge et al., 2023). Zebrafish were maintained at 28 °C in a water 59 recirculating system under a 12 h light: dark photoperiod. The pH level of the water is maintained at 7.0 – 8.0 and conductivity at 800 μs/cm (Lawrence, 2007). Adult were fed 4 times a day alternating Zebrafeed (400-600 μm, Sparos, Portugal) and *Artemia nauplii.* Mating was performed using a single couple tank 0.8 L capacity and eggs were collected, washed with egg water, and incubated at 28 °C until hatching. At 10 dpf (days post fertilization) zebrafish were introduced into the recirculating system. Larvae were fed from 5 dpf with Zebrafeed (<100 μm). During growth, the size of the feed was increased and *Artemia nauplii* was introduced (from 10 dpf).

The in vivo study consisted of two complementary experimental designs aimed at investigating the effects of chronic and acute β-glucan supplementation.

### Chronic BGs supplementation

Starting from the second week of life, fish were fed until 27 weeks post hatching (wph) with Zebrafeed (400–600 μm) either as control diet or supplemented with 1,3-1,6 β-glucans (BGs) at two concentrations: 12.5 mg/kg BW/day and 125 mg/kg BW/day. The fish of voluntary feed intake (FI) was estimated at 50 mg/day, corresponding to an FI ratio of 5% BW/day. Based on these values, diets were supplemented to provide approximately 12.5 mg/kg BW/day and 125 mg/kg BW/day of BGs, corresponding to 0.25 g/kg feed and 2.5 g/kg feed, respectively.

These doses (12.5 mg/kg BW and 125 mg/kg BW) were selected based on previous studies demonstrating immunomodulatory effects of β-glucans in fish species (Bagni et al., 2000; Misra et al., 2005; Fronte et al., 2019; Ching et al., 2020). Number animals at the end of supplementation:

- n = 10 control animals
- n = 9 animals (12.5 mg/kg BW)
- n = 10 animals (125 mg/kg BW)

This experimental approach was used to evaluate the effects of BG supplementation on survival and age-associated phenotypes. Survival was monitored longitudinally and analyzed using Kaplan–Meier survival analysis, with n ≥ 40 animals per group. At defined ages, organs including brain, liver, kidney, muscle and heart were collected for histological and molecular analyses.

### Acute BG intervention

*Nothobranchius furzeri* were fed with Zebrafeed fortified with 1.25 mg/g of 1,3-1,6 β-glucans (corresponding to a dose of 125 mg/kg BW, assuming a daily food intake of 100 mg per fish) for a week starting at age 26 weeks.

Fish received BG-fortified feed at:

- 0 mg/kg BW (n = 5)
- 125 mg/kg BW (n = 5)

At the end of treatment, organs were collected for histological and immunofluorescence analyses.

### Inclusion of 1,3-1,6 β-glucans in the food

MacroGard^®^ (Biorigin^©^, São Paulo, Brazil) was used as source of 1,3-1,6 β-glucans, derived from *Saccharomyces cerevisiae* cell wall). The 1,3-1,6 β-glucans feed inclusion was performed as described by Fronte et al. (2019), to reach a final dose of 12.5 mg/kg BW and 125 mg/kg BW.

### Tissue processing and histology

Collected organs were processed for histological and immunofluorescence analyses.

#### Lipofuscin detection

Unstained slides were mounted using a mounting solution for lipofuscin detection. As lipofuscin is autofluorescent, no staining is required to produce its emission when it is stimulated at 488 nm. Via the use of the apotome microscope, lipofuscin was obtained.

#### Histology

According to the established protocols, slides were stained with haematoxylin and eosin (H&E) and Sirius Red.

#### Protocol for H&E staining

Immerse the sections in Hematoxalin for 3 minutes (min). The sections are rinsed with water for 5 min. Then, the sections are stained with Eosin for 45 seconds. The sections are rinsed with water for 5 min. Subsequently, the sections were dehydrated at different concentrations of ethanol (3 x 5’ 95% ethanol, 3 x 5’ 100% ethanol, 2 x 10’ xylene. The sections were mounted using Permount (xylene based).

#### Sirius Red Fast Green protocol

Sirius red and fast green staining (9046, Chondrex, Inc.) was used to detect muscle collagen deposits. Collagen fibers appeared magenta, while the non-collagen proteins were green. Slides were incubated with 0.3 ml Dye Solution for 30 min. After, the Dye Solution was aspirated, and the slides were rinsed until the water runs clean. The slides were dehydrated and covered with coverslip.

#### Hepatocyte vacuolization

Hepatocellular vacuolation would be considered as lipidosis or steatosis. Cytoplasmic inclusions of lipids are found in hepatocytes. Fish tend to accumulate more glycogen/lipids, in fact have larger vacuoles than mammals. The high hepatocellular vacuolation in *Nfu* is a natural phenomenon (De Cicco et al., 2009; Godoy, R. S. *et al*. 2019). Histologically we observe cytoplasmic inclusions of lipid in hepatocyte. In agreement, with De Cicco, I semi-quantitatively measured hepatocytes vacuolization by defining 4 different degrees (De Cicco et al., 2009). In addition, we classified the vacuoles, into glycogen-like or lipid vacuoles. The lipid type manifested itself as unstained cytoplasmic vesicles with sharp edges. The glycogen-like type was characterized by irregular vacuoles containing slightly flocculent material (Zac, et al., 2020).

#### Kidney and muscle analysis

Dilation of the renal tubules is a typical condition of aging. I measured the diameter of each individual renal tubule using 3 images/section by 3 sections. Moreover, I calculated the presence of precipitate in renal tubules and necrotic tubules Necrotic renal tubules show sloughing of tubular cells into the lumen and severe tubular leakage. We measured the increase in thickness of Bowman’s capsule relative to the area of the glomerulus. The renal corpuscle is formed by a centrally located glomerulus surrounded peripherally by a Bowman’s capsule. To quantify the increment of in thickness of Bowman’s capsule I calculated the ratio between the thickness of Bowman’s capsule (pixels2) compared to the total area (pixels2) of the renal corpuscle (Bowman’s capsule and Glomerulus).

#### Feret’s size

HE-stained muscle fibers were individually measured via Feret’s size from imageJ. Feret’s size measure diameters derived from the distance of two tangents to the fiber boundary in a well-defined orientation. All fibers from 3 images for 3 sections for each animal were analyzed.

#### Fibrosis

Collagen deposition was used to measure fibrosis in the muscle. Collagen deposition was quantified analyzing the percentage area of the magenta fibers (collagen). The percentage area of the image was measured by setting the same threshold for all images.

### Organotypic *ex-vivo* brain slice cultures

To investigate the direct effects of BGs on brain tissue independently of systemic influences, organotypic brain slice cultures were established.

Brains were isolated from 5-, 10- and 27-week-post-hatching fish.

Experiments are performed under sterile conditions with tools sterilised by autoclaving prior to use. 70% EtOH is used to sterilize equipment and surfaces throughout the experiment.

1. The fish were euthanized in MS-222 (200 mg/L), and placed in 0.05% NaOH for 30 seconds, followed by 70% EtOH for 30 seconds. This is done to sterilize the body and prevent the brain from becoming contaminated with bacteria during dissection.
2. The fish were decapitated using scissors.
3. The heads were transferred in the dissecting medium and brains were dissected using a stereomicroscope. The longer it takes to extract the brain, the less likely it is that the slices will survive.
4. The brains extracted were placed it on the micrometric slide and, using a precision micro-knife, cut it in half sagittally.
5. The cultured brain slices were extracted and appropriately positioned on the semipermeable membranes (PICM0RG50, Merk Life Science, Germany) containing culture medium, place the six well plate into the incubator set at a temperature of 28°C and 5% CO2.

Slices were cultured at 28 °C and 5% CO₂ in DMEM/F12 medium supplemented with:

- 10% fetal calf serum
- 10% sterile water
- 0.033% insulin
- 511 μM ascorbic acid
- 1% penicillin/streptomycin
- glucose adjusted to 0.4%

Culture medium was replaced daily.

The slices were treated with 1,3-1,6 β-glucans (BGs) at 8 mg / L concentration (Brogi et al., 2021) in culture medium for three days to observe the effects of 1,3-1,6 β-glucans in autophagy at different aging. Moreover, the cultured brain slices extracted of Nothobranchius of 5 wph and Tg(mpeg1.1.: EGFP+) zebrafish were also treated with 100 nM of Bafilomycin A1 for 3 days to observe if 1,3-1,6 β-glucans is inductor of autophagy and restore autophagy in condition of impairment of this mechanism.

#### Pharmacological manipulation of autophagy and microglia

Brain slices were treated for 3 days with:

- β-glucans (BGs): 8 mg/L
- Bafilomycin A1: 10 nM or 100 nM
- plx5622: 20 μM

BG stock solution (MacroGard®) was prepared in sterile water and sonicated for 2 min (30 s ON/OFF cycles).

Bafilomycin A1, an inhibitor of the vacuolar H⁺-ATPase, dissolved in DMSO and diluted to final concentrations in culture medium at 100 nM.

plx5622, a selective CSF1R inhibitor that induces microglial depletion, was dissolved in DMSO and diluted to final concentrations in culture medium at 20 µM.

To investigate the effects of BGs directly on brain, *Nfu* were treated with 8 mg/L of BGs for three days at different ages (5, 10, and 27 wph). Following treatment, tissues were collected and analyzed by immunofluorescence and proteomic analyses. To assess autophagic flux, fish at 5 wph were exposed for three days to BGs, Bafilomycin A1 (100 nM), or their combination. Four experimental groups were analyzed: control, BGs, Bafilomycin A1, and BGs + Bafilomycin A1. Autophagic flux was further evaluated at 27 wph using CYTO-ID staining. In this experiment, fish were treated for three days with BGs alone or BGs in combination with Bafilomycin A1 (100 nM). To evaluate the role of microglia, fish were treated for three days with plx5622 (20 μM), a CSF1R inhibitor known to induce microglial depletion. Microglial depletion was confirmed at 20 wph, and subsequent experiments included four groups: control, BGs, plx5622, and BGs + plx5622.

Throughout the study, the number of biological replicates (n) refers to the number of half brains analyzed.

#### Lysotracker and MitoTracker staining

First of all, the use of Lysotracker (LysoTracker™ Green DND-26, special packaging; L7526; Thermo Fisher) and MitoTracker in the ex-vivo model was validated, and the optimal incubation parameters were determined by monitoring dye penetration every 5 min (Lysotracker) or every 30 min (MitoTracker). Not fixed cultured brain slices (at 27 wph) were stained with Lysotracker, marker of lysosomes, or MitoTracker, marker of Mitochondria. Lysotracker was introduced to the medium at the concentration of 75 μM for 45 min at 28 ° C in an incubator. Then, 3 washes of 10 min with medium were acquire using a Zeiss LSM 900 with Airyscan. The lyophilised dried powder of MitoTracker reagent was reconstituted in DMSO, as described in the Product Datasheet. The MitoTracker was added at the concentration of 500 nM in medium for 90 min at 28 ° C and 5% CO2 in an incubator, followed by three 10-min washes. The cultured brain slices were fixed in PFA 4% as for immunofluorescence analyses and they are viewed using confocal Zeiss LSM 900 with Airyscan.

#### CYTO-ID Autophagic detection

CYTO-ID® Autophagy detection kit 2.0 (ENZ-KIT175, Enzo) in the *ex-vivo* model was validated, given that this kit is designed for cell cultures. Then, 2 μL of CYTO-ID® Green Detection Reagent and 1 μL of Hoechst 33342 Nuclear Stain were added to the medium of the cultured brain slices. Initially, every 30 min it was observed whether the Reagent had penetrated the cultured brain slices in confocal instrument. 2 μL of CYTO-ID® Green Detection Reagent 2 for 90 min at 28 ° C and 5% CO2 in the incubator are the best parameters for the action of this kit. The cultured brain slices were rinse three times for 10 min and were fixed in PFA 4% as for immunofluorescence analyses and they are viewed using Confocal (Stellaris 5, Leica).

### Human iPSC-derived cortical neurons

Culture plates were coated ON with Poly-D-Lysine (50 μg/mL, Sigma-Aldrich) at 37 °C in a humidified incubator (5% CO₂). The following day, plates were washed and coated ON with Mouse Laminin I (10 μg/mL, Merck Millipore) under the same conditions. Human iPSC-derived cortical neurons were thawed at 37 °C, diluted 1:5 in HBSS (Thermo Fisher Scientific) and centrifuged at 200 g for 4 min at room temperature. The cell pellet was resuspended in WiBiTi maintenance medium supplemented with 2 μM Y-27632 (ROCK inhibitor) and 20 μM DAPT (Notch inhibitor) and plated at a density of 200,000 cells/cm² on laminin-coated plates. Neurons were cultured from 10 to 90 days in vitro (DIV) under different maintenance media depending on the stage of differentiation. From 10–15 DIV, neurons were maintained in WiBiTi medium, consisting of DMEM/F-12 supplemented with 2 mM glutamine, 1 mM sodium pyruvate, 1 mM non-essential amino acids, 0.05 mM β-mercaptoethanol, 10 μM 53AH, 10 μM LDN193189 hydrochloride, 1 μM RepSox, N-2 supplement and B-27 supplement minus vitamin A. From 16–32 DIV, neurons were maintained in N2B27 medium, composed of DMEM/F-12 supplemented with glutamine (2 mM), sodium pyruvate (1 mM), non-essential amino acids (1 mM), β-mercaptoethanol (0.05 mM), N-2 supplement and B-27 supplement minus vitamin A.Between 32–70 DIV, neurons were cultured in Neurobasal Young medium, consisting of Neurobasal medium supplemented with 2 mM glutamine, 1 mM sodium pyruvate, 0.05 mM β-mercaptoethanol, N-2 supplement, B-27 supplement minus vitamin A, 0.5 mM ascorbic acid and recombinant human BDNF (20 ng/mL). From 70–90 DIV, neurons were maintained in Neurobasal Old medium, identical to the Young medium except that B-27 supplement replaced B-27 minus vitamin A. Culture medium was partially refreshed every 2–3 days. Neurons were passaged every two weeks. Cells were washed with Versene, detached with Acutase for 10 min at 37 °C, diluted 1:5 in PBS, and centrifuged at 200 g for 4 min at room temperature. The pellet was resuspended in the appropriate maintenance medium supplemented with 2 μM Y-27632, 20 μM DAPT and 10 μM LDN193189 hydrochloride. After 24 h, the medium was replaced with fresh unsupplemented medium. The final passage was performed on 24-well glass-bottom plates for confocal imaging following immunofluorescence staining.

#### β-glucan and Bafilomycin A1 treatments in iPSC-derived neurons

After maintenance of iPSC-derived human cortical neurons up to 87 (or 89) days DIV, cells were treated with BGs, 8 mg/L and Bafilomycin A1 (10 or 100 nM). For the acute treatment (24 h) experiment four experimental groups were analyzed: control, BGs (8 mg/L), Bafilomycin A1 (100 nM), and Bafilomycin A1 (100 nM) + BGs (8 mg/L). For the long treatment (72 h) experiment, six experimental groups were included: control, BGs (8 mg/L), Bafilomycin A1 (10 nM), Bafilomycin A1 (10 nM) + BGs (8 mg/L), Bafilomycin A1 (100 nM), and Bafilomycin A1 (100 nM) + BGs (8 mg/L).

### Immunofluorescence

#### Immunofluorescence staining on tissue slides and Proteostat™ Aggresome staining

After washing in PBS, tissue slides were subjected to antigen retrieval using citrate buffer (2.94 g trisodium citrate dihydrate, 0.05% Tween-20 in H₂O, pH 6.0). The solution was heated to 95 °C, and slides were immersed for 2 min, repeating the cycle three times.

For aggresome staining, sections were incubated with ProteoStat™ Aggresome Detection Kit (Enzo Life Sciences) at 1:2000 in PBS for 3 min, rinsed in PBS, de-stained in 1% acetic acid for 40 min, and rinsed in PBS.

To prevent non-specific staining, sections were blocked with 5% BSA and 0.3% Triton X-100 in PBS for 2 h at room temperature. Slides were incubated ON at 4 °C with primary antibodies (Table 2) diluted in 1% BSA and 0.1% Triton X-100 in PBS. After washing (3 × 5 min PBS), samples were incubated with secondary antibodies (1:500) for 2 h at room temperature. Sections were washed again and stained with DAPI (10 μg/ml) for 10 min, rinse in PBS for 10 min and mounted with Fluoroshield mounting medium (Sigma-Aldrich).

**Table 2:** Primary antibodies used in this study for immunofluorescence experiments.

| Antibody Primary | Marker of | Working Solution | Product type | Code |
| --- | --- | --- | --- | --- |
| **GFAP** | **Gliosis** | **1:400** | **Rabbit Polyclonal** | **GTX128741, Genetex** |
| **LAMP1** | **Lysosomes** | **1:500** | **Rabbit Polyclonal** | **ab24170, Abcam** |
| **LC3B** | **Autophagy** | **1:500** | **Rabbit Polyclonal** | **ab48394, Abcam** |
| **LCP1** | **Inflammation** | **1:200** | **Rabbit Polyclonal** | **GTX124420, Genetex** |
| **Nitrotyrosine** | **Protein peroxidation** | **1:500** | **Rabbit Polyclonal** | **A-21285, Thermo Fisher** |
| **βIII-Tubulin** | **Neurons** | **1:10000** | **Mouse Monoclonal** | **801201, BioLegend** |

#### Immunofluorescence staining of *ex-vivo* brain slices

Brain slices were fixed with 4% PFA for 1 hour (h) and washed in PBS(3 x 15 min). Samples were blocked with 5% BSA and 0.1% Triton X-100 in PBS for 4 h at room temperature under gentle agitation. Slices were incubated with primary antibodies (Table 2) diluted in 1% BSA and 0.1% Triton X-100 in PBS for 3 days at 4 °C under agitation. After primary antibody incubation, slices were washed in PBS (3 × 15 min) and incubated with secondary antibodies ON. Samples were then washed (3 × 15 min), stained with DAPI (10 μg/ml) ON, and washed again in PBS (3 × 15 min). For optical clearing, slices were incubated in A2 Scale solution (10% glycerol, 4 M urea, 0.1% Triton X-100) until transparent (up to 1 week). Fluorescence preservation was achieved using S4D25(0) Scale solution (40% sorbitol, 10% glycerol, 4 M urea, 25% DMSO).

#### Immunofluorescence staining of iPSC-derived human neurons

Human iPSC-derived cortical neurons (DIV 90) were fixed with 2% PFA in PBS for 15 min at room temperature and washed three times in PBS. Cells were permeabilized and blocked in 10% goat serum and 0.5% Triton X-100 in PBS for 1 h at room temperature. Samples were incubated ON at 4 °C with primary antibodies (Table 2) diluted in 10% goat serum in PBS. After washing (3 × 10 min PBS), cells were incubated with secondary antibodies (1:1000) for 1 h at room temperature. After secondary antibody incubation, neurons were washed in PBS (3 × 10 min), stained with DAPI (10 μg/ml), washed in PBS (1 × 15 min), and mounted with Aqua-Poly/Mount (Polysciences). Slides were allowed to cure for 72 h before confocal imaging.

### Microscopy

Histological images were acquired in bright field using a Nikon Eclipse E400 microscope. Fluorescence images were acquired using a Zeiss Apotome.2 microscope equipped with an Axiocam 807 mono camera, and images were processed using the ZEN Blue software suite. Z-stack images were acquired using 40× and 63× oil immersion objectives, with a z-step of 0.5 μm. For each sample, three areas were acquired from three sections. Cultured brain slices were imaged using a Zeiss LSM 900 confocal microscope with Airyscan 2 using a 63× oil immersion objective in Airyscan mode, acquiring z-stacks of 10 focal planes with a step size of 2.5 μm. For Tg(mpeg1.1:EGFP) brain slice cultures, z-stack images were acquired using a Leica Stellaris 5 confocal microscope with a 63× objective (NA 1.28), using a 2 μm step size over a 20 μm thickness. Brain slices treated with plx5622 were imaged under the same conditions with a 2.5 μm step size spanning 25 μm thickness. For each experiment, six areas were acquired for 3–5 cultured brain slices, depending on the experimental condition. iPSC-derived human neurons were imaged using a Leica Stellaris 5 confocal microscope with a 63× objective (NA 1.28), acquiring 10 z-stacks with a step size of 0.5 μm. For each group, three wells (24-well plates) were analyzed, acquiring 10 images per well.

### Image quantification and sample size

Images were quantified using Fiji (ImageJ) by measuring the percentage of fluorescent area relative to the total image area (pixels²). The same threshold was applied to all images within each experiment. For the chronic diet experiment, three groups were compared (Control, 12.5 mg/kg BW, and 125 mg/kg BW), including all animals surviving to 27 wph (Control *n*=10, 12.5 mg/kg BW *n*=9, 125 mg/kg BW *n*=10). For the acute diet experiment, Control and 125 mg/kg BW groups were analyzed (*n*=5 per group). For each phenotype, three images per section from three sections were analyzed (9 images per sample). For brain slice cultures, 3–5 samples per group were used depending on the experiment. Specifically, n=5 per group were used for age-dependent autophagy and Bafilomycin A1 experiments, as well as Tg(mpeg1.1:EGFP) brain slice cultures. CYTO-ID, LysoTracker, MitoTracker, and LCP1 analyses included n=4 samples per group, whereas plx5622-treated brain slice cultures included n=3 samples per group. For each culture, five images were analyzed. For iPSC-derived neurons, three wells per condition (24-well plates) were analyzed, acquiring 10 images per well. Neuronal survival and LC3 puncta were quantified from six z-stacks spaced by 0.5 μm per image. Thresholds for fluorescence quantification were set as follows: Nitrotyrosine (150–255 cardiac sections; 100–255 hepatic sections), GFAP (75–255 brain), LC3 (110–255 liver; 120–255 brain; 100–255 muscle), NDUFS1 (180–255), and lipofuscin (90–255 liver; 85–255 brain; 100–255 heart). Lysosomal activity was calculated as the ratio between aggregate area and LAMP1-positive lysosomal area (thresholds: LAMP1 100–255; aggresomes 90–255). Inflammation was quantified by manually counting LCP1-positive cells relative to total cells. For 3D analyses of LC3, LysoTracker, MitoTracker, and CYTO-ID, z-stack images were analyzed using the 3D Object Counter plugin in Fiji. Images were converted to 8-bit, and identical thresholds and size filters (≥1 pixel) were applied across samples (LC3: 25–255; LysoTracker: 50–255; MitoTracker: 30–255; CYTO-ID: 20–255). Quantified parameters included number of objects, object volume, percentage area normalized to cell area, and integrated density. Microglial morphology in Tg(mpeg1.1:EGFP) fish was quantified by measuring cell sphericity after 3D surface reconstruction. LC3 puncta in iPSC-derived neurons were quantified as LC3-positive objects within the βIII-tubulin–defined neuronal surface (Surface and Spot tools; thresholds: βIII-tubulin 11–60, LC3 2.5–60) and expressed as dots per neuron, while neuronal survival was assessed by counting βIII-tubulin–positive neurons.

### Proteomics analyses

#### Preparation of cultured brain slices for

After 3 days of cultivation the cultured brain slices were collected and twice for one min in fresh 1% PBS with 1% phosphatase and protease inhibitors. Subsequently, they were weighed in the precision balance. The cultured brain slices were collected and snap-frozen in liquid nitrogen and stored at -80 ° C.

#### Sample preparation for proteomics analysis

Tissues (cultured brain slices) were resuspended in PBS to a concentration of 200 μg/μl cells before adding 2x lysis buffer (final concentration of 5% SDS, 100 mM HEPES and 50 mM DTT). The samples were sonicated (Bioruptor Plus, Diagenode, Belgium) for 10 cycles (30 sec ON/60 sec OFF) at a high setting at 20°C, followed by boiling at 95°C for 5 min. Reduction was followed by alkylation with iodoacetamide (final concentration 15 mM) for 30 min at room temperature in the dark. Samples were acidified with phosphoric acid (final concentration 2.5%), and seven times the sample volume of S-trap binding buffer was added (100 mM TEAB, 90% methanol). Samples were bound on S-trap micro spin columns (Protifi) and washed three times with binding buffer. Trypsin in 50 mM TEAB pH 7.55 was added to the samples (1 μg per sample) and incubated for 1 h at 47°C. The samples were eluted in three steps with 50 mM TEAB pH 7.55, elution buffer 1 (0.2% formic acid in water) and elution buffer 2 (50% acetonitrile and 0.2% formic acid). The eluates were dried using a speed vacuum centrifuge (Eppendorf Concentrator Plus, Eppendorf AG, Germany). The samples were resuspended in Evosep buffer A (0.1% formic acid in water) and sonicated (Bioruptor Plus, Diagenode, Belgium) for 3 cycles (60 sec ON/30 sec OFF) at a high setting at 20°C. The samples were loaded on Evotips (Evosep) according to the manufacturer’s instructions. In short, Evotips were first washed with Evosep buffer B (acetonitrile, 0.1% formic acid), conditioned with 100% isopropanol and equilibrated with Evosep buffer A. Afterwards, the samples were loaded on the Evotips and washed with Evosep buffer A. The loaded Evotips were topped up with buffer A and stored until the measurement.

#### LC-MS Data independent analysis (DIA)

Peptides were separated using the Evosep One system (Evosep, Odense, Denmark) equipped with a 15 cm x 150 μm i.d. packed with a 1.9 μm Reprosil-Pur C18 bead column (Evosep Endurance, EV-1106, PepSep, Marslev, Denmark). The samples were run with a pre-programmed proprietary Evosep gradient of 44 min (30 samples per day) using water and 0.1% formic acid and solvent B acetonitrile and 0.1% formic acid as solvents. The LC was coupled to an Orbitrap Exploris 480 (Thermo Fisher Scientific, Bremen, Germany) using PepSep Sprayers and a Proxeon nanospray source. The peptides were introduced into the mass spectrometer via a PepSep Emitter 360-μm outer diameter × 20-μm inner diameter, heated at 300°C, and a spray voltage of 2.2 kV was applied. The injection capillary temperature was set at 300°C. The radio frequency ion funnel was set to 30%. For DIA data acquisition, full scan mass spectrometry (MS) spectra with a mass range of 350–1650 m/z were acquired in profile mode in the Orbitrap with a resolution of 120,000 FWHM. The default charge state was set to 2+, and the filling time was set at a maximum of 60 ms with a limitation of 3 × 106 ions. DIA scans were acquired with 40 mass window segments of differing widths across the MS1 mass range. Higher collisional dissociation fragmentation (stepped normalized collision energy; 25, 27.5, and 30%) was applied, and MS/MS spectra were acquired with a resolution of 30,000 FWHM with a fixed first mass of 200 m/z after accumulation of 1 × 106 ions or after filling time of 45 ms (whichever occurred first). Data were acquired in profile mode. For data acquisition and processing of the raw data, Xcalibur 4.4 (Thermo) and Tune version 3.1 were used.

#### Proteomic data processing

DIA raw data were analysed using the directDIA pipeline in Spectronaut (v.16.2, Biognosysis, Switzerland). The data were searched against an in-house species-specific database (*Nothobranchius furzeri* 59.154 entries) and contaminant (247 entries) SwissProt database. The data were searched with the following variable modifications: oxidation (M) and acetyl (protein N-term). A maximum of 2 missed cleavages for trypsin and 5 variable modifications were allowed. The identifications were filtered to satisfy an FDR of 1% at the peptide and protein levels. Relative quantification was performed in Spectronaut for each paired comparison using the replicate samples from each condition. The data were exported, and further data analyses and visualization were performed with R studio using in-house pipelines and scripts.

### Data Analysis of proteomics data

#### Identification of DEPs

DEPs were identified with |logFC| >0 and qvalue <0.10. The DEPs with logFC <0 or logFC <0 were considered downregulated or upregulated genes, respectively. DEPs common among datasets were screened using the Venn diagram software (Venny 2.0; https://bioinfogp.cnb.csic.es/tools/venny/index2.0.2.html).

#### Protein-protein interaction network construction and module analysis

Protein–protein interaction (PPI) networks were generated using the STRING database (http://string-db.org). Clustering was performed within STRING using the identified protein set as background, and Cellular Component enrichment and PPI enrichment statistics were used to identify functional modules.

### Principal component analysis (PCA) and heatmap analysis

PCA of their hierarchical clustering were performed using the ComplexHeatmap 2.0.0, pcaMethods 1.76.0, and ggpubr 0.2 packages of R.

### Statistics Analysis

GraphPad Prism 8 program (GraphPad, San Diego, CA, USA) and ImageJ software were used to perform statistical analysis. All the data were expressed as Mean ± standard error of the mean (S.E.M). Data regarding survival curves were subjected to the log-rank test. Data regarding percentage of vacuole’s type were calculated with Chi^2^ test. Some data were analysed by One-way ANOVA trend analysis test. Other data were analysed by Two-way ANOVA test with Tukey’s multiple comparison test to compare the different groups. Other data were analysed by Student’s t-test calculated adjusted p-value with FDR. Differences between treatments were considered significant for *p* ≤ 0.05 (*), *p* ≤ 0.01 (**), *p* ≤ 0.001 (***) and *p* ≤ 0.00001 (****).

## ACKNOWLEDGEMENTS

The authors gratefully acknowledge Dr. Cinzia Caterino (Leibniz Institute on Aging – Fritz Lipmann Institute, Jena) for technical support. We thank the Proteomics Facility at the Leibniz Institute on Aging – Fritz Lipmann Institute for proteomic analyses, with special thanks to Dr. Emilio Cirri for his support. The Tg(mpeg1.1) transgenic line was kindly provided by Prof. Ambrosio (Biogem, Ariano Irpino, Italy). We also thank Filippo Maria Santorelli for insightful discussions, valuable suggestions, and for hosting visits to his laboratory.

## DATA AVAILABILITY STATEMENT

The protein database generated in this study has been deposited in MassIVE (UCSD) and is publicly available at: http://massive.ucsd.edu/ProteoSAFe/status.jsp?task=0180a0273c254c2da0e7cbe8a56841bd.

## AUTHOR CONTRIBUTIONS

L.B. conceived the study and contributed to the conceptualization, designed and performed the experiments, carried out formal analysis and data curation, and wrote the original draft of the manuscript.

B.F. and A.C. contributed to the conceptualization of the study.

M.M. contributed to the conceptualization of the study, provided scientific supervision, and critically revised the manuscript.

A.C. supervised the project and critically revised the manuscript.

F.T. and F.C. provided human neuronal cultures and contributed to experimental support and scientific discussion.

All authors reviewed and approved the final version of the manuscript.

