## Supplementary Figures and caption for "1,3-1,6 β-glucans reduce aging hallmarks in multiple organs and rapidly induce mitochondrial biogenesis and autophagy via direct effect on the killifish brain and human neurons"

**Caption Supplementary Figures**

**Figure S1. β-glucan treatment modulates body weight, lipofuscin accumulation, and autophagy across multiple tissues.** (A) Body weight (BW) analysis of animals subjected to chronic β-glucans (BGs) treatment. (B) Body weight analyzed separately for male and female animals across experimental groups. (C) Representative fluorescence images showing lipofuscin autofluorescence (green) in liver sections from control animals and fish treated with BGs (12.5 mg/kg BW and 125 mg/kg BW). Arrowheads indicate lipofuscin granules. Scale bar: 10 µm. (D) Quantification of lipofuscin-positive area (% area) in liver. (E) Representative fluorescence images showing lipofuscin autofluorescence in heart tissue from control and BG-treated animals. Arrowheads indicate lipofuscin deposits. Scale bar: 5 µm. (F) Quantification of lipofuscin-positive area (% area) in heart. (G) Representative confocal images of LC3 immunostaining (green) with DAPI (blue) in liver tissue, showing autophagosome distribution in control and BG-treated animals. Scale bar: 10 µm. (H) Quantification of LC3-positive area (% area) in liver. (I) Representative confocal images of LC3 immunostaining (green) with DAPI (blue) in muscle tissue. Scale bar: 10 µm. (J) Quantification of LC3-positive area (% area) in muscle. Data are presented as mean ± SEM. Statistical significance was determined using one-way ANOVA followed by Tukey's multiple comparison test. Sample sizes: $n = 10$ control, $n = 9$ (12.5 mg/kg BW), $n = 10$ (125 mg/kg BW). *$p < 0.05$, **$p < 0.01$, ***$p < 0.001$, ****$p < 0.0001$.

**Figure S2. β-glucan treatment modulates lysosomal activity, microglial markers and oxidative stress across multiple tissues.** (A) Representative confocal images showing LAMP1 (green), aggresome staining (red), and DAPI (blue) in liver sections from control animals and fish treated with β-glucans (BGs). Scale bar: 10 µm. (B) Quantification of aggresome-positive area relative to LAMP1-positive area. (C) Representative confocal images of LCP1 immunostaining (red) with DAPI (blue) in gut sections from control and BG-treated animals. Scale bar: 10 µm. (D) Quantification of LCP1+/total cells in gut tissue. (E) Representative confocal images showing nitrotyrosine staining (green) with DAPI (blue) in liver tissue. Scale bar: 10 µm. (F) Quantification of nitrotyrosine-positive area (% area) in liver. (G) Representative confocal images showing nitrotyrosine staining (green) with DAPI (blue) in heart tissue from control and BG-treated animals. Scale bar: 10 µm. (H) Quantification of nitrotyrosine-positive area (% area) in heart. Data are presented as mean ± SEM. Statistical significance was determined using one-way ANOVA followed by Tukey's multiple comparison test. Sample sizes: $n = 10$ control, $n = 9$ (12.5 mg/kg BW), $n = 10$ (125 mg/kg BW). *$p < 0.05$, **$p < 0.01$, ***$p < 0.001$, ****$p < 0.0001$.

**Figure S3. β-glucan treatment does not induce major structural alterations in skeletal muscle and liver tissues.** (A) Representative hematoxylin and eosin (H&E) staining of skeletal muscle sections from control animals and fish treated with β-glucans (BGs). Scale bar: 20 µm. (B) Quantification of Feret's diameter in skeletal muscle sections. (C) Representative Sirius Red staining of skeletal muscle sections highlighting collagen deposition. Scale bar: 20 µm. (D) Quantification of collagen-positive area (% area) measured from Sirius Red staining. (E) Representative hematoxylin and eosin (H&E) staining of liver sections showing hepatocyte morphology and vacuolization. Scale bar: 50 µm. (F) Quantification of hepatocyte vacuolization in liver sections. (G) Quantification of vacuole composition in hepatocytes, distinguishing glycogen-like and lipid vacuoles. Data are presented as mean ± SEM. Statistical significance was determined using one-way ANOVA followed by Tukey's multiple comparison test. Chi-square ($\chi^2$) test was used to analyze vacuole distribution.

Sample sizes: n = 10 control, n = 9 (12.5 mg/kg BW), n = 10 (125 mg/kg BW). *p < 0.05, **p < 0.01, ***p < 0.001.

**Figure S4. β-glucan treatment does not significantly alter kidney morphology.** (A) Representative hematoxylin and eosin (H&E) staining of kidney sections from control animals and fish treated with β-glucans (BGs; 12.5 mg/kg BW and 125 mg/kg BW). Scale bar: 50 µm. (B) Quantification of Bowman's capsule enlargement. (C) Quantification of % renal tubules with precipitates. (D) Quantification of % necrotic renal tubules. (E) Quantification of renal tubular area ($pixels^2$). Data are presented as mean ± SEM. Statistical significance was determined using one-way ANOVA followed by Tukey's multiple comparison test. Sample sizes: n = 10 control, n = 9 (12.5 mg/kg BW), n = 10 (125 mg/kg BW). *p < 0.05, **p < 0.01, ***p < 0.001.

**Figure S5. Microglia depletion reduces lysosomal and autophagic markers in aged *ex-vivo* brain.** (A) Representative confocal images of Cyto-ID staining (green) in aged *ex-vivo* brain (27 weeks post hatching or wph) treated with Bafilomycin A1 alone or in combination with β-glucans (BGs). Nuclei are counterstained with DAPI (blue). Scale bar: 10 µm. (B) Quantification of autophagosome number per cell. (C) Quantification of autophagosome volume ($pixel^3$). (D) Quantification of autophagosome area ($pixel^2$/cell). (E) Representative confocal images of LCP1 immunostaining (green) with DAPI (blue) in *ex-vivo* brain (20 wph) under control conditions or treated with plx5622 (20 µM) for 3 days. Scale bar: 10 µm. (F) Quantification of LCP1+/total cells showing efficient microglial depletion following plx5622 treatment. Data are presented as mean ± SEM. Statistical analysis was performed using unpaired two-tailed Student's t-test or one-way ANOVA followed by Tukey's multiple comparison test, as indicated. Sample sizes: n = 4 for each group. *p < 0.05, ***p < 0.001.

**Figure S6. Protein interaction network of proteins upregulated by BGs treatment at 10 wph.** (A) Protein–protein interaction network of proteins commonly upregulated during normal aging in vivo and downregulated by β-glucans (BGs) treatment *ex-vivo*. Functional clustering highlights pathways related to metabolic pathways (red nodes). (B) Protein–protein interaction network of proteins downregulated by BG treatment at 27 wph *ex-vivo*. Functional clustering highlights pathways related to synapses (red nodes), ATP binding (blue nodes), and spectrin-associated cytoskeletal organization (green nodes). Network visualization was generated using protein–protein interaction databases and pathway enrichment analysis.

**Figure S7. KEGG pathway enrichment analysis of proteins upregulated by BGs treatment in aged brain *ex-vivo*.** (A) KEGG pathway map of oxidative phosphorylation showing proteins upregulated following β-glucan (BG) treatment *ex-vivo* in aged brains (27 wph), identified by proteomic analysis. Highlighted proteins belong to mitochondrial respiratory chain complexes, indicating modulation of mitochondrial metabolism. (B) KEGG pathway map of the lysosome pathway, showing differential regulation of proteins involved in lysosomal membrane components, lysosomal enzymes, and vesicular transport machinery, suggesting modulation of lysosomal function and autophagy-related processes by BG treatment in aging *ex-vivo*. (C) KEGG pathway map of the GABAergic synapse pathway highlighting proteins involved in synaptic signaling, vesicle trafficking, and receptor regulation, indicating that BG treatment affects neuronal synaptic pathways in the aging brain *ex-vivo*. Differentially expressed proteins are highlighted within each pathway map according to their relative abundance change (color scale from green to red indicating down- to up-regulation). Pathway visualization was generated using KEGG pathway analysis and Pathview mapping tools.

**Figure S8. Proteomic network analysis reveals pathways modulated by BGs treatment in the adult brain.** (A) Protein–protein interaction network of proteins upregulated following β-glucans

(BGs) treatment *ex-vivo* in adult brains (10 wph), identified by proteomic analysis. Functional clustering highlights pathways related to synaptic processes (red nodes) and axon-associated proteins (blue nodes). Network visualization was generated using protein–protein interaction databases and pathway enrichment analysis.

**Figure S9. BGs treatment partially rescues neuronal survival following autophagy inhibition in iPSC-derived human cortical neurons.** (A) Representative confocal images of iPSC-derived human cortical neurons stained for βIII-tubulin (red) to visualize neuronal morphology. Nuclei are counterstained with DAPI (blue). Neuronal cultures were treated with 100 nM Bafilomycin A1 for 24 hours (h) or 72 h, either alone or in combination with β-glucans (BGs) treatment. Representative conditions shown include Bafilomycin A1 (72 h), Bafilomycin A1 + BGs (72 h), Bafilomycin A1 (24 h), and Bafilomycin A1 + BGs (24 h). Scale bar: 25 μm. (B) Quantification of neuronal survival, calculated as the ratio of βIII-tubulin–positive neurons to total DAPI-positive nuclei, used as a measure of neuronal viability across experimental conditions. Data are presented as mean ± SEM. Statistical analysis was performed using two-way ANOVA followed by Tukey's multiple comparison test. Sample sizes are indicated in the graphs (number of independent neuronal cultures analyzed per condition). BAF 100 = 100 nM Bafilomycin A1. $*p < 0.05$, $**p < 0.01$, $***p < 0.001$, $****p < 0.0001$.

**Figure S10. β-glucan treatment modulates LC3 accumulation in iPSC-derived human cortical neurons following autophagy inhibition.** (A) Representative confocal images of iPSC-derived human cortical neurons stained for βIII-tubulin (red) to visualize neuronal morphology and LC3 (green) to detect autophagosomes. Nuclei are counterstained with DAPI (blue). Neurons were analyzed after 72 h of treatment under the following conditions: control, β-glucan (BGs), Bafilomycin A1 (10 nM), Bafilomycin A1 (100 nM), BGs + Bafilomycin A1 (10 nM), and BGs + Bafilomycin A1 (100 nM). LC3 signal was analyzed exclusively in βIII-tubulin–positive neurons. Scale bar: 25 μm. (B) Quantification of the number of autophagosomes normalized to βIII-tubulin area ($\mu m^2$), showing the effects of BGs and autophagy inhibition with Bafilomycin A1 (10 nM and 100 nM) after 72 h. Data are presented as mean ± SEM. Statistical analysis was performed using two-way ANOVA followed by Tukey's multiple comparison test. Sample sizes are indicated in the graph. BAF = Bafilomycin A1; BAF 10 = 10 nM Bafilomycin A1; BAF 100 = 100 nM Bafilomycin A1. $*p < 0.05$, $**p < 0.01$, $***p < 0.001$.
